# Pathology of dose dependent inocula of H5N8 avian influenza virus in experimentally infected chicken

**DOI:** 10.64898/2026.01.27.700741

**Authors:** Boopathi Ponnusamy, Manoj Kumar, Harshad Vinayakrao Murugkar, Shanmugasundaram Nagarajan, Chakradhar Tosh, Sivasankar Panickan, Dhruv Desai, Semmannan Kalaiyarasu, Dhanapal Senthilkumar, Katherukamem Rajukumar, Siddharth Gautam, Vijendra Pal Singh, Aniket Sanyal

## Abstract

In the present study, we assessed the pathogenicity of H5N8 avian influenza viruses belongs to the clade 2.3.4.4b in chicken. Birds of three different dose groups, 10^2^, 10^4^, and 10^6^ EID_50_ were used in the study. No mortality was observed in 10^2^ EID0 group. Percent cumulative mortality of 10^4^ and 10^6^ EID_50_ group was 66.67 and 100 %, respectively. Varying duration of MDT of 3.2 and 2 days was observed in 10^4^ and 10^6^ EID_50_ group, respectively. The CID_50_ of virus was found to be 10^4.5^ EID_50_. High no. of viral RNA copies were found both in oropharyngeal and cloacal swabs and in various organs of birds infected in 10^4^ and 10^6^ EID_50_ group. Significant gross and histological changes and presence of viral antigen in various organs were observed in 10^4^ and 10^6^ EID_50_ group. So, the study concludes that Indian HPAI, H5N8 isolates are highly pathogenic in nature to chicken by affecting most organs systemically. CID_50_ of this H5N8 virus indicates poor adaption in chicken and it implies poor transmission possibility of this virus for host species in field condition. Though this virus is highly pathogenic in nature as that of HPAI, H5N1 viruses, absence of endothelial staining in most organ attributes variation in replication process and pathogenesis from HPAI, H5N1 viruses. Hence, further studies need to be done to elucidate the pathobiology of this virus in various bird species.

**Highlights:**

- H5N8 virus belong to the clade 2.3.4.4b, Indian isolate is highly pathogenic in nature as that of HPAIV, H5N1.
- The dose inocula, 10^2^ EID_50_ is noninfectious to chicken.
- The dose inocula, 10^4^ and 10^6^ EID_50_ had caused significant mortality in the inoculated chicken with MDT of 2 and 3.2 days, respectively.
- H5N8 virus was detected with high viral titres in clocal and oral shedding and in multiple organ with the dose inocula, 10^4^ and 10^6^ EID_50_.
- 10^4^ and 10^6^ EID_50_ of H5N8 inocula virus caused significant gross and histological changes in multiple organs and viral antigens were detected in respective organs.

## 1. Introduction

Avian influenza is caused by type A, H5 and H7 influenza viruses of genus influenza virus A and family orthomyxoviridae. Influenza viruses are pleomorphic and enveloped RNA viruses which are separated into 8 segments, code for 10-18 proteins. Further, these viruses are divided into different subtypes based on the antigenicity of their envelope surface glycoproteins, i.e. haemagglutinin (HA) (H1 to H18) and neuraminidase (NA) (N1 to N11) [1–3]. All naturally occurring high pathogenicity avian influenza (HPAI) strains isolated to date have been either of the H5 or H7 subtype, with a subset of H5 or H7 isolates being of low pathogenicity [4]. Further, high pathogenicity subtypes cause acute clinical diseases and heavy economic losses in chickens, turkeys, and other bird species population [4]. Aquatic bird species, anseriformes (ducks, geese, swans) and Charadriiformes (shorebirds, gulls, alcids), are the main source of influenza virus infecting domestic avian species and mammals [5,6]. H5N1 is the representative subtype of HPAIV in Asia and these viruses have been discretely involved in reassortment events and resulting in generation of H5 strains with novel NA subtypes or donated gene segments to other co circulating strains [7–10]. Because of Extensive genetic divergence and reassortment between other subtypes of influenza viruses, these HPAI viruses with an H5 subtype continue to undergo substantial evolution [11]. Further, HPAI H5 subtype viruses belong to the lineage of Goose/ Guangdong (Gs/GD) has spread across worldwide and causing outbreaks in domestic and wild bird species [12]. Since 2003, A/goose/Guandong/1/96–like viruses contributed hemagglutinin (HA) gene to influenza A(H5N1) viruses have become endemic in Bangladesh, China, Egypt, India, Indonesia, and Vietnam [13]. So, highly mutating and reassortant nature of these viruses paved the way for outbreak of HPAI viruses with H5 subtype with various NA subtype, N1, N2,N3,N5,N6, ad N8 in different continents, Asia, Europe, and North america [3, 14–17]. As an event of various outbreak of Gs/GD in different continents, novel HPAI virus, H5N8 was reported in two zoos of india also on October, 2016 (Nagarajan 2017). The reported isolates were from delhi National Zoological Park, Delhi, and Gandhi Zoological Park, Gwalior, Madhya Pradesh [18]. Phylogentic analysis revealed that this viruses belong to the clade 2.3.4.4b and different 7:1 reassortants of H5N8 viruses isolated in May 2016 from wild birds in the Russian Federation and China [18]. In continuation to this, HPAI, H5N8 virus outbreaks were reported in domestic chicken and duck from different states of india, caused substantial mortality and economic loss [19–21]. Thus, it is imperative to study the pathobiology of this novel viruses, H5N8, belong to the clade 2.3.4.4 to know the replication in various organs and its pathogenesis. In field conditions, pathogenicity and transmission of HPAIVs could be affected by various factors such as host species, the age of the susceptible population, the viral dose, biosecurity measures followed, and breeding condition [22, 23]. Though reports available about pathogenicity [24–27] and or pathobiology [28–31] of HPAI, H5N8 viruses, no data is available pertaining to Indian HPAI, H5N8 isolates. Further, reports are scanty regarding the dose dependent inoculation and outcome of infection and mortality in chickens. Hence, the present study was carried out to investigate pathobiology of 2016 Indian isolate, HPAI H5N8 virus with different dose inocula in the chicken model.

## 2. Materials and methods

### 2.1 Virus

Highly pathogenic avian influenza, H5N8 virus A/chicken/India/11CA01/2016 belong to the clade 2.3.4.4b was used in this experiment. The virus was obtained in the form of amnio-allantoic fluid (AAF), from “Avian influenza virus repository” of ICAR-National Institute of High Security Animal Diseases (NIHSAD), Bhopal. Haemagglutination of virus was done before determining the infectivity [32].

### 2.2 Determination of infectivity of stock virus

Specific Pathogen Free (SPF) embryonated chicken eggs (ECEs) were used to determine infectivity of the virus. The eggs were procured from the SPF Unit of ICAR-NIHSAD and SPF Division of Venkateshwara Hatcheries Pvt. Limited, Pune, India. The infectivity of the stock virus was determined by calculating 50% egg infective dose (EID50) in 9-11 days old embryonated chicken eggs. The EID50 was calculated as per the standard method described [33].

### 2.3 Animals and housing

Specific Pathogen Free (SPF) chickens were procured from ICAR-NIHSAD SPF Unit for the experiment. Before inoculation of virus, serum samples were collected from representative number of chickens to confirm that the birds were serologically negative for avain influenza virus (AIV) by haemagglutinin inhibition (HI) test [32]. Further, oral and cloacal swabs were collected and virus shedding was determined by quantitative real-time RT-PCR (qRRT-PCR) [34] to ensure the absence of AIV. Each experimental group was housed separately in negative pressure isolators with inlet and exhausted HEPA-filtered ventilation within the animal bio-containment facility (ABSL3) of ICAR-NIHSAD. Ad libidum feed and water access was provided to all birds. All the study related protocols or procedures were reviewed and approved by the Institute Animal Ethics Committee (IAEC approval No. 61/IAEC/ HSADL/12).

### 2.4 Experimental design and sampling

To evaluate the pathogenicity of virus, H5N8 virus A/chicken/India/11CA01/2016, 72 number of SPF chickens of either sex aged 4-6 weeks were used for intranasal inoculation. Inoculation was done after 48 hours (hrs) of acclimatization period. Three different groups of birds (n = 18 / group) were infected with three different doses of virus, 10^2^, 10^4^, and 10^6^ EID_50_ in the volume of 100 ul. These three groups are hereafter referred with the dose inocula being used in the study viz., 10^2^, 10^4^, and 10^6^ EID_50_ groups. Sham infected control (n=18) was inoculated with sterile allantoic fliud. From each group, 3 birds were scheduled for humane sacrifice (cervical dislocation) at the time intervals, 6, 12, 24, and 48 hours post infection (hpi) for tissue collection. Remaining 6 birds of all groups were decided to subject for observation up to 14 days followed by sacrifice on the same day. Mean Death Time (MDT) was calculated by using the formula (MDT = Σ (dxM)/N, Where, d - ith day M - Number of birds died on ith day, N - Total number of birds used for the study) and 50 % Chicken Infectious Dose (CID_50_) calculation was done as per standard protocol [33] to determine the infective dose of H5N8 virus. Surviving birds at the end of experiment will be subjected for serological examination, HI, to know the presence or absence of seroconversion [32]. Oralphayngeal and cloacal swabs were collected in 650 ul of sterile Phosphate Buffered Saline (PBS) after 24 hpi till 14 days post infection (dpi) for the viral RNA quantification by qRT-PCR from all groups [34]. Only, birds showing positive titre were subjected for virus RNA quanitification calculation. Tissue samples, nasal cavity, trachea, lungs, heart, liver, spleen, intestines, pancreas were collected in RNA Later solution for viral RNA copy number quantification by qRRT-PCR [34]. In addition to the aforementioned tissues, thymus and bursa were collected for histopathology (HP) and immunohistochemistry (IHC) in 4 % Paraformaldehyde (PFA).

### 2.5 ViralRNA quantification for swabs and tissue extractions

Oropharyngeal and cloacal swabs were subjected for qRRT-PCR to determine viral load after infection. Viral RNA was extracted from the processed swabs using QIAamp Viral RNA mini kit (Qiagen, Germany Cat.No. 52904) as per the recommended protocol of the manufacturer. For extraction of viral RNA from tissues, 50-100 mg of tissue was taken for RNA extraction. Ten percent homogenous suspension (w/v) in 1X pre-chilled PBS was prepared and homogenised tissue suspension was used for RNA extraction by organic method using TRizol® reagent (Ambion, Life Technologies, USA, Cat.No.15596-018). In vitro transcribed matrix gene RNA using T7 polymerase in Roche IVT RNA synthesis (Roche, Germany), obtained from the clone repository, ICAR-NIHSAD was used for standard curve preparation. Prior to dilution, quantity of RNA transcripts was determined by Qubit® Fluorometer using QuantiT™ RNA Assay Kit (Invitrogen, USA) as per manufacturer’s protocol. Copy numbers were calculated by using the formula (Concentration of RNA in grams /μl / [length of amplicon Χ 340]) Χ 6.022 Χ 1023 = Number of molecules / μl (http://www.qiagen.com/resources/info/guidelines_ rtpcr/quantifying.aspx). Tenfold serial dilutions of the quantified RNA standards of matrix gene were prepared in TE buffer and 1 μl of each dilution was used as template for preparation of standard curve using primers for individual IVT RNAs. This qRRT-PCR was carried out in Light Cycler® 480 Real-Time PCR System II (Roche, USA) using Super Script III Platinum One-Step qRT-PCR Kit (Invitrogen, USA, Cat. No. 11732020). The primers used are GAAGGARTRCCTGAGTCTA (forward - 5’→3’) and YRTCGTCARCATCCACAG (Reverse - 5’→3’). The sequence of probe is TGCTGTTCCTKYCGRTAYTCTT, which is labelled with FAM and BHQ-1. By using same matrix gene primers and probe, reaction was performed to know the virus quantification in swabs and tissues. Pre-quantified 10^-4^ dilution of IVT RNA was used in each test run as positive control. From this viral RNA copy numbers of sample were calculated by using obtained Ct values.

### 2.6 Histopathology (HP) and immunohistochemistry (IHC)

Tissue samples collected and fixed in 4 % paraformaldehyde were embedded with paraffin, sectioned at the thickness, 4-6 μm, and stained with hematoxylin and eosin (H&E) [35]. Chicken polyclonal sera against the clade, 2.3.4.4b H5N8 viruses was used for immunhistochemical detection of virus antigen in tissues, as per the previously published protocol [36]. The chicken polyclonal sera were obtained from the serum repository of ICAR-NIHSAD.

### 2.6. Data analysis

Data were presented as mean ± standard deviation of mean, which was analyzed by t-test (Welchon) and ordinary one way and two-way ANOVA using GraphPad prism software version 10.4.1 (San Diego, CA, USA). Results were considered statistically significant, if P-values were *≤0.05 or **≤0.01 or ***≤0.001or ****<0.0001.

## 3. Results

### 3.1. Clinical signs, Mortality, Mean death time and Chicken infectious doe (CID_50_)

No clinical signs and mortaity mortality were observed in mock infected and 10^2^ EID_50_ group. After the 48 hpi interval, remaining 6 birds were kept till the end of experiment i.e. upto 14 dpi and euthanised on 14 dpi, were found to be sero-negative to H5 virus. In 10^4^ and 10^6^ EID_50_ infected groups, birds started showing clinical signs from 24- and 48-hour post infection (hpi), respectively. The predominant clinical signs include dullness, huddling, anorexia, prostration, dropping of eyelids and the nervous signs viz., torticollis and circling in sacrificed and found dead birds. Nervous signs were more pronounced in found dead birds (53-132 hpi) of 10^4^ EID50 group than 10^6^ EID_50_ group. In 10^4^ and 10^6^ EID_50_ groups, mortality was observed as presented in the table. 1. Their survival rate is depicted in figure. 1. In 10^4^ EID_50_ group, after 48 hpi interval, four out of six birds succumbed to the infection between 53 −132 hpi. Two surviving birds euthanised on 14 dpi were found seronegative against H5 antibodies by HI test. In 10^6^ EID_50_ group, after 24 hpi interval, the mortality started around 45 hpi and all birds were dead by 47 hpi. MDT and CID_50_ were calculated based on exact death timing of birds (Table.1, below).

**Fig. 1:**
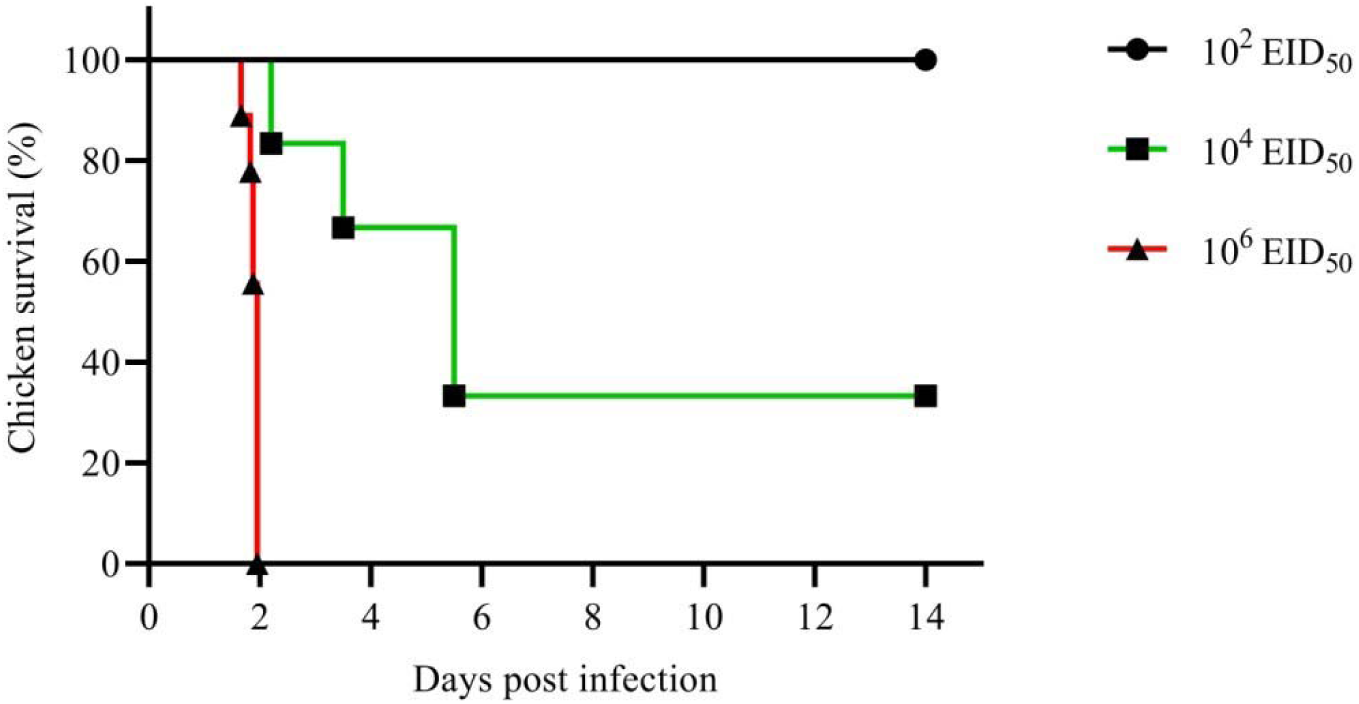
Survival of chickens after infected intranasally with 10^2^, 10^4^, and 10^6^ EID_50_ of H5N8 (A/chicken/India/11CA01/2016) virus.

**Table 1:**
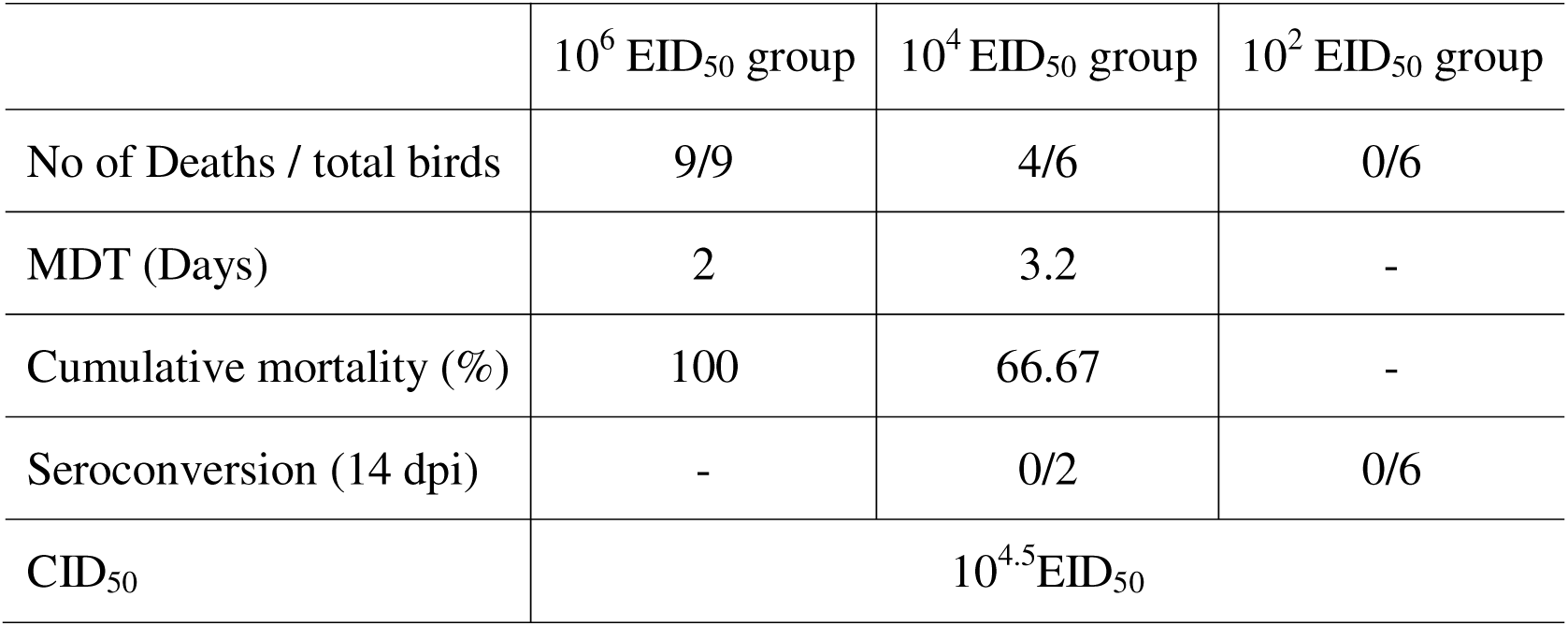
Cumulative mortality (%), MDT, and CID50 of HPAI, H5N8 virus inoculated chickens with different EID_50_.

| | $10^6$ EID<sub>50</sub> group | $10^4$ EID<sub>50</sub> group | $10^2$ EID<sub>50</sub> group |
| --- | --- | --- | --- |
| No of Deaths / total birds | 9/9 | 4/6 | 0/6 |
| MDT (Days) | 2 | 3.2 | - |
| Cumulative mortality (%) | 100 | 66.67 | - |
| Seroconversion (14 dpi) | - | 0/2 | 0/6 |
| CID<sub>50</sub> | $10^{4.5}$ EID<sub>50</sub> | | |

Percent Cumulative Mortality from each group was derived and mentioned (Table.1). The MDT of 10^4^ EID_50_ and 10^6^ EID_50_ group was 3.16 days and 2 days, respectively. On other hand, experimental challenge of HPAIV, H5N8 with 10^2^ EID_50_ dose inocula did not cause any mortality in chicken. In our study, the resultant CID_50_ of A/chicken/India/11CA01/2016 (H5N8) virus was 10^4.5^ EID_50_.

### 3.2. Viral RNA quantification in oropharyngeal and clocal shedding

In mock infected and 10^2^ EID_50_ group, all the oropharyngeal and cloacal swabs screened through 1 to 14 dpi were negative for viral RNA. In 10^4^ and 10^6^ EID_50_ group viral RNA was detected both in oral and cloacal swabs and the copy numbers are depicted in Fig. 2. In 10^4^ EID_50_ group viral RNA copies were detected from 1 to 5 dpi (Fig. 2). After 5 dpi, 2 surviving birds did not shed the detectable amount of viral RNA in both oropharyngeal and cloacal excretions till 14 dpi. Highest viral RNA copies were detected at 4 and 3 dpi in oral (2.8×10^8^) and cloacal (1×10^8^) swabs, respectively (Fig. 2). In 10^6^ EID_50_ group, viral RNA copies were detected at 1 dpi only, as no birds survived after that period (Fig. 2). Both in 104 and 106 EID50 group, oral shedding was higher than cloacal shedding from 1-5 dpi except 3 dpi cloacal shedding of 104 EID50 group (Fig. 2). Between the groups, oral shedding of 106 EID_50_ group (6.5×10^7^) was significantly higher than 104 EID_50_ group. In 10^4^ EID_50_ group, at 3 and 4 dpi, cloacal (1×10^8^) and oral (2.8×10^8^) shedding of H5N8 virus was significantly higher than other dpi viral shedding.

**Fig. 2:**
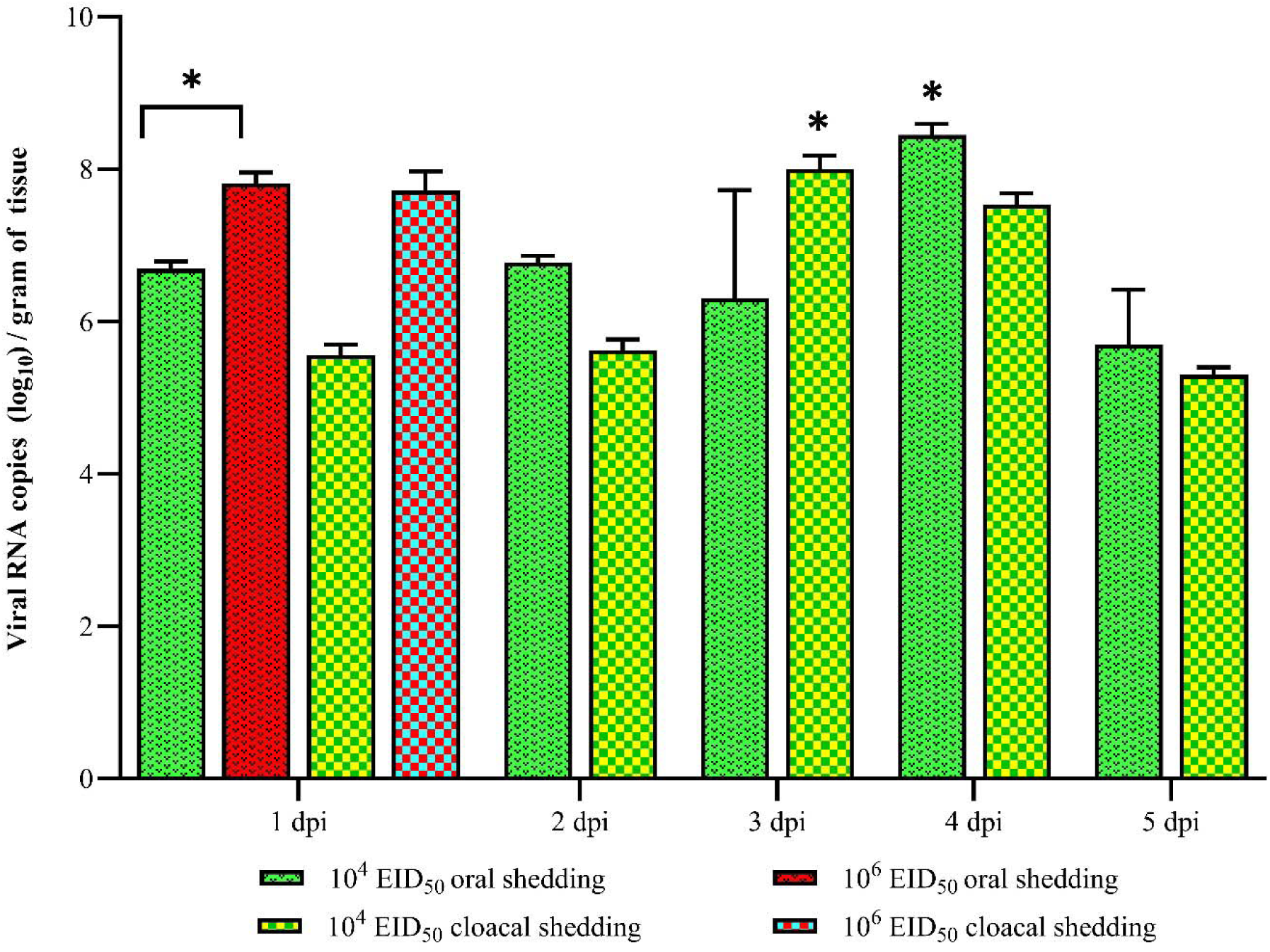
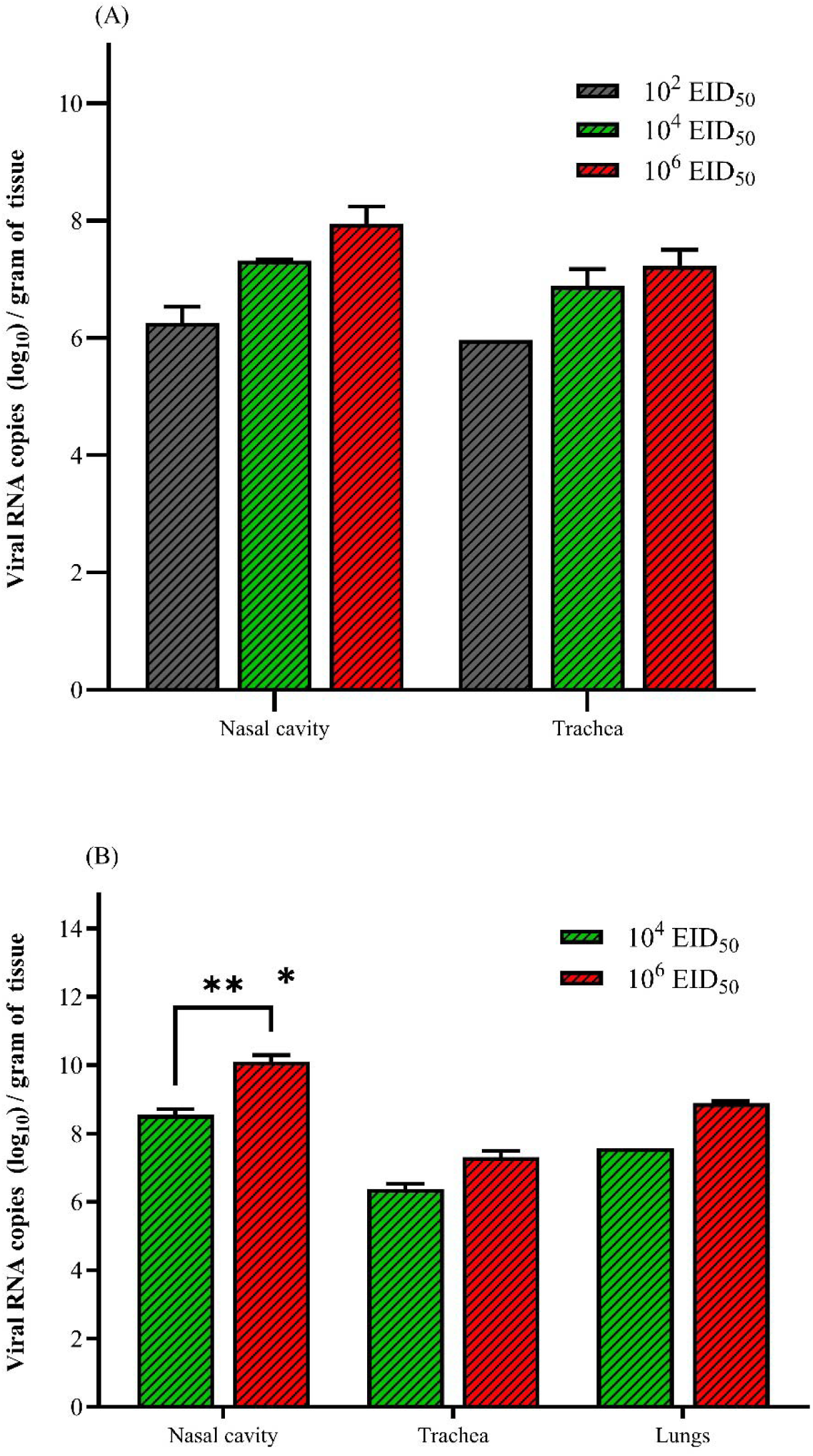
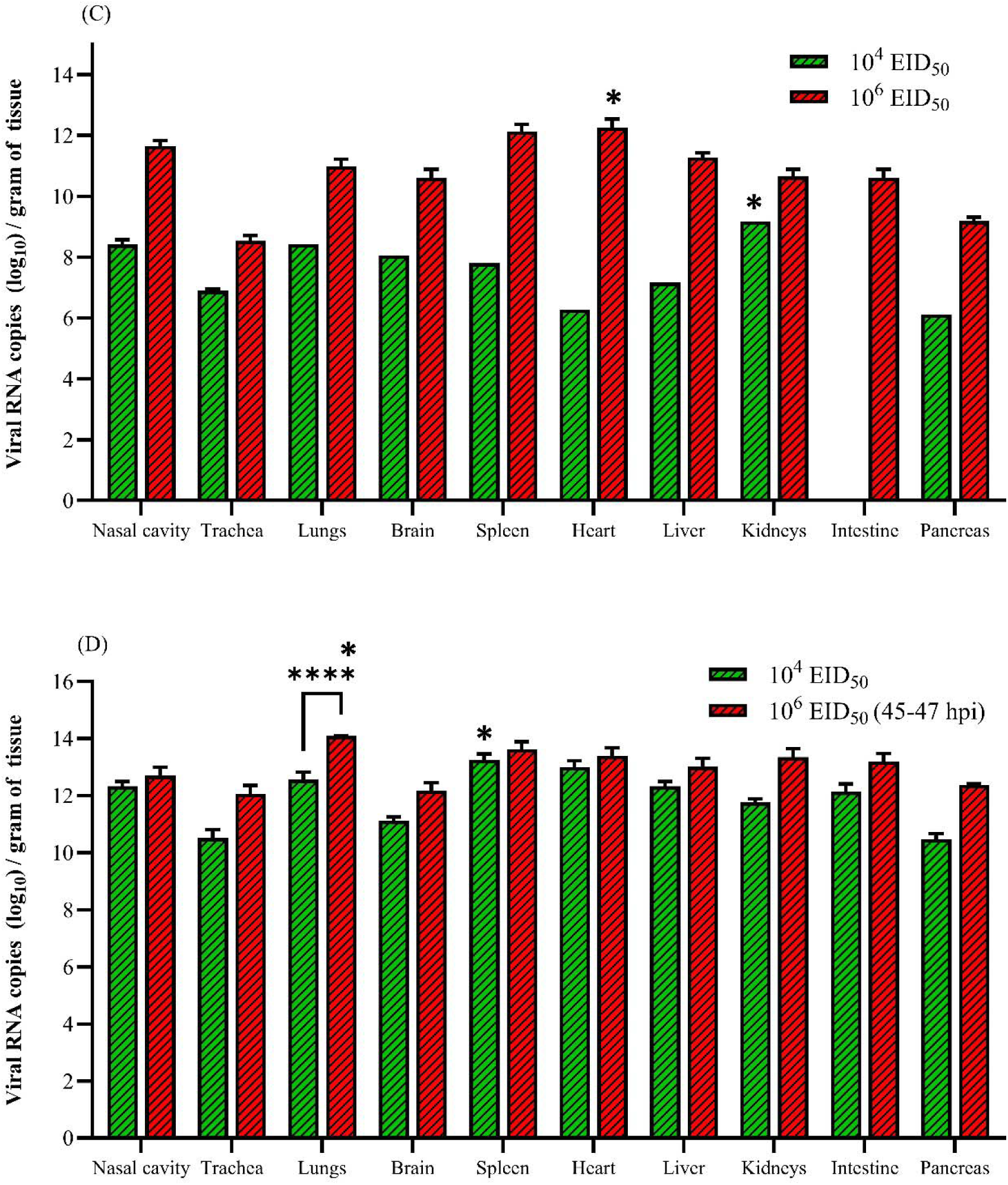
Viral RNA copies in oral and cloacal shedding of chickens infected with 10^4^ and 10^6^ EID_50_ of HPAIV, H5N8 from 1 to 5 dpi. Data are expressed as Mean ± SD and analyzed by two-way ANNOVA to know the significance between the groups and the significance are shown by asterisk mark with lines. Further, the significance of oral and cloacal shedding within the group is shown by asterisk without lines by the one-way ANOVA. n=3 in treatment groups, 10^4^ and 10^6^ EID_50_. Differences are considered statistically significant, if P-values were *≤0.05.

### 3.3. Virus RNA quantification in various systemic organs

The viral RNA copy numbers in H5N8 infected chicken varied between different organs at different intervals (Figs. 3A-E) of each group and in found dead birds. At 6 hpi, viral RNA was detected in upper respiratory tract viz., trachea and nasal cavity in all the three dose groups (Fig.3A). No significant difference was observed within and as well as between the groups. After 6 hpi, viral RNA was not detected in any organ in all the tested intervals in 10^2^ EID_50_ group till 14 dpi. At 12 hpi, virus was detected in trachea, nasal cavity, and lungs in 10^4^ and 10^6^ EID_50_ dose groups (Fig 3). However, in 10^4^ EID_50_ dose group, only one out of three birds showed viral RNA in lungs at 12 hpi (Fig. 3B). No significant difference was observed within and as well as between the groups. At 24 hpi, virus was systemically present in all organs in 10^6^ EID_50_ dose group (Fig 3C). However, in 104 EID50 dose group, only one bird showed systemic presence of virus in all screened organs except intestine and other two birds showed viral RNA copies in nasal cavity, trachea and lungs only (Fig. 3C). Kidneys (1.52×10^9^) in 10^4^ EID_50_ group was observed with significantly higher viral RNA copies than all other organs except brain. In 10^6^ EID_50_ group, heart (1.81×10^12^) was observed with significantly higher viral RNA copies than the organs, trachea and pancreas. Between 45-47 hpi, all remaining 9 chickens were succumbed by H5N8 infection in 10^6^ EID_50_ group. So, data of 3 out of 9 deceased chickens were compared (Fig. 3D) with mean viral RNA copies of 48 hpi euthanized chickens of 10^4^ EID_50_ group. At this time point, in 10^4^ EID_50_, group, spleen (1.79×10^13^) was observed with significantly higher viral RNA copies than all other organs except heart. In 10^6^ EID_50_ group, lungs (1.23×10^14^) were observed with significantly higher viral RNA copies than all the other organs. Moreover, between the groups, lungs (1.23×10^14^) of 10^6^ EID_50_ group were observed with significantly higher viral RNA copies than the treatment group, 10^4^ EID_50_. As no data is available from 10^6^ EID_50_ dose group after 48 hpi, viral RNA detected in 10^4^ EID_50_ dose group at 53-132 hpi was presented alone (Fig. 3E). After 48 hpi, in 10^4^ EID_50_ dose group, four out of six birds succumbed to infection (53-132 hpi) had high viral RNA copies in all tested organs (Fig. 3E). Remaining two survived chickens were observed with negative for viral RNA copies in all the tested organs.

**Fig. 3:**
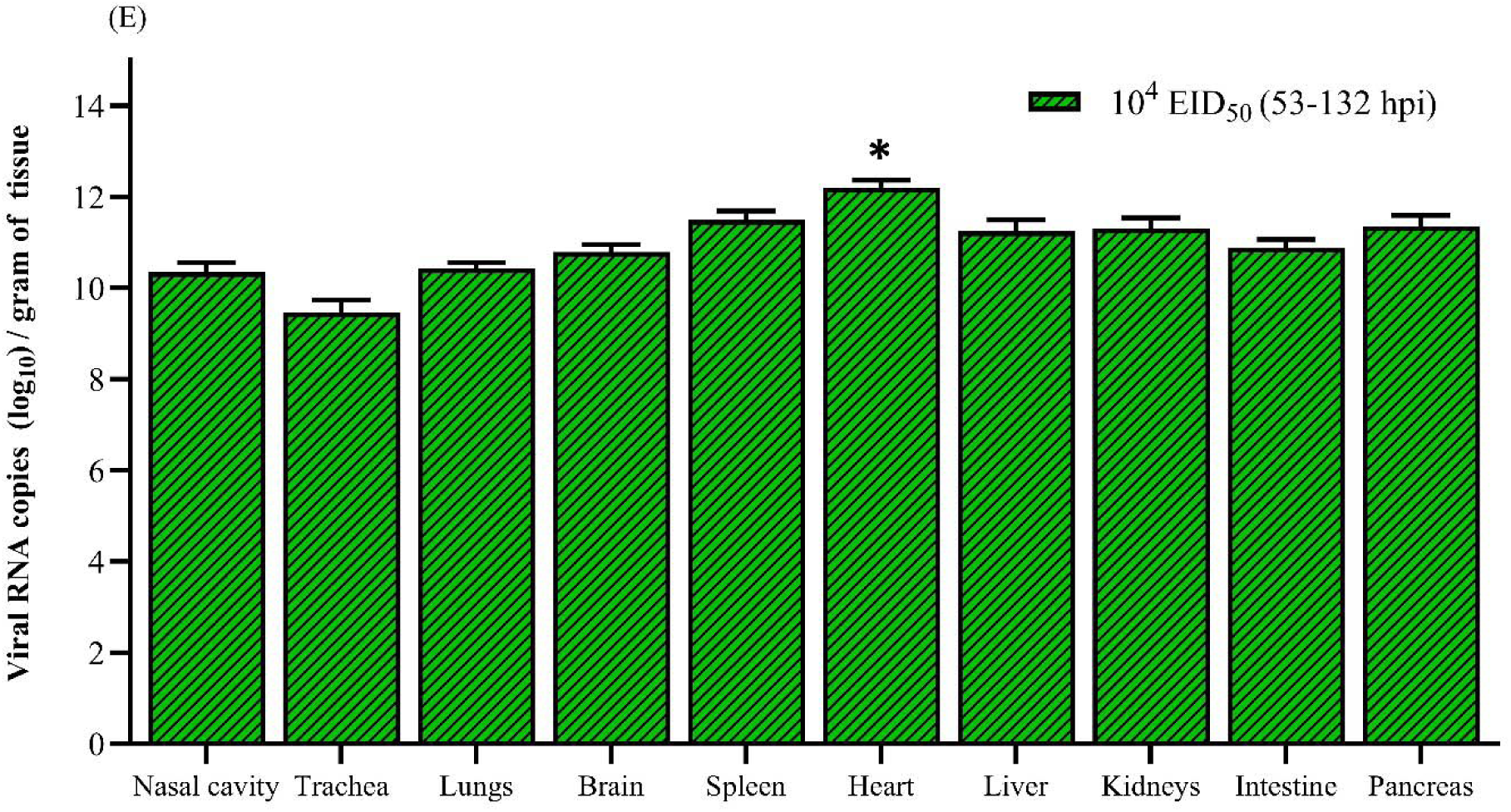
Viral RNA copies in different organs of chickens infected with 10^2^, 10^4^, and 10^6^ EID_50_ of HPAIV, H5N8 at 6 hpi (A), 12 hpi (B), 24 hpi (C), 48 hpi (D), and > 48 hpi (E). Data are expressed as Mean ± SD and analysed by two-way ANNOVA to know the significance between the groups and the significance are shown by asterisk mark with lines. Further, at all time lines except at 6 hpi, the significance between various organs within the group is shown by asterisk without lines by the one-way ANOVA. At 6 hpi, data are analysed by T test-Welchon (Two tailed). The error bar represents standard error of mean. n=3 and 1 with and without error bars, respectively at 6, 12, 24 and 48 hpi in treatment groups. n = 4 in 10^4^ EID_50_ group after 48 hpi (53-132 hpi). Differences are considered statistically significant, if P-values were *≤0.05 or **≤0.01 or ****<0.0001.

### 3.4. Gross and histopathological findings

All birds in the mock infected and 10^2^ EID_50_ group did not show any gross pathological lesions and the organs were normal in shape, size, colour and consistency. In 10^4^ and 10^6^ EID_50_ group, various organs showed significant gross changes at different hpi (Supplementary Table. 1 & Figs. 4-6). In lungs (Fig. 4), congestion, haemorrahge and oedema were significantly noticed. In addition, congestion of trachea and meninges of brain, pale liver, parenchymal mottling and whitish foci of spleen (Fig. 5), fluid filled pericardial sac of heart, subcutaneous and thigh muscle haemorrhages (Fig 6), and thymic regression were observed in affected birds. Other organs did not show appreciable gross lesions.

**Fig. 4:**
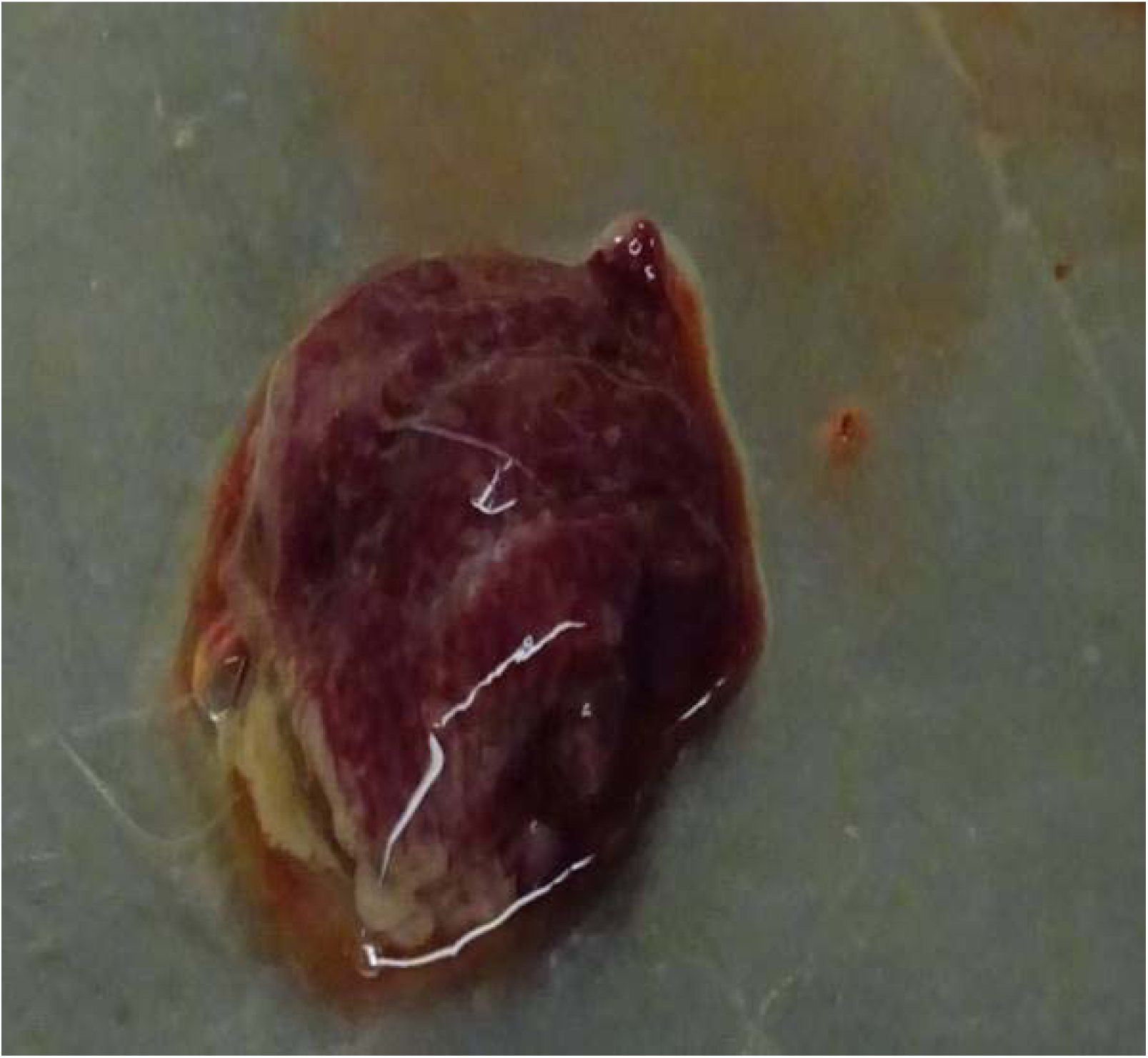
10^6^ EID_50_ group - dead bird (47 hpi) - Lung: Showing severe congestion, oedema and haemmorrhage.

**Fig. 5:**
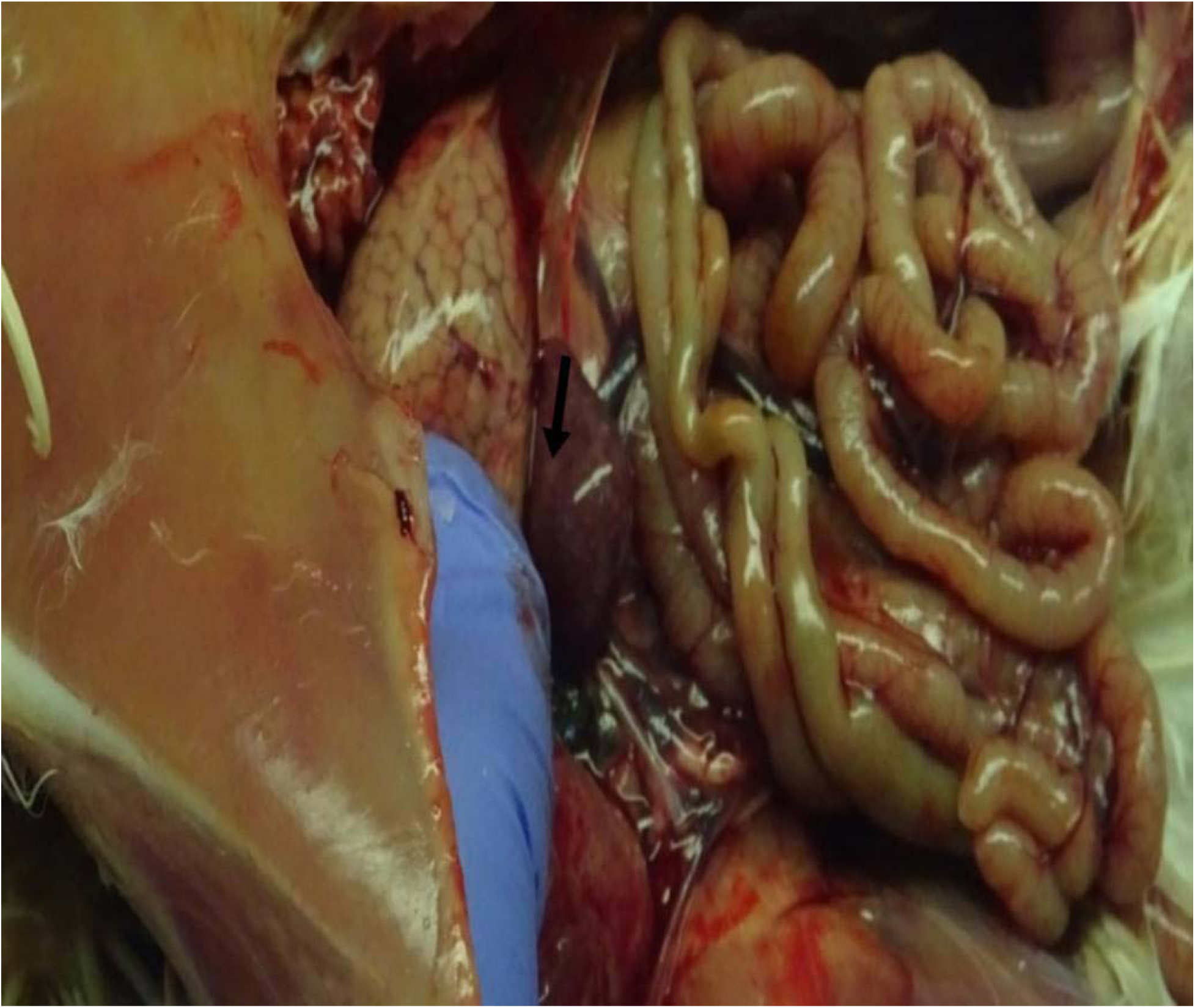
10^6^ EID_50_ group - dead bird (47 hpi) - Spleen: Showing irregular size of greyish - white foci on surface (arrow).

**Fig. 6:**
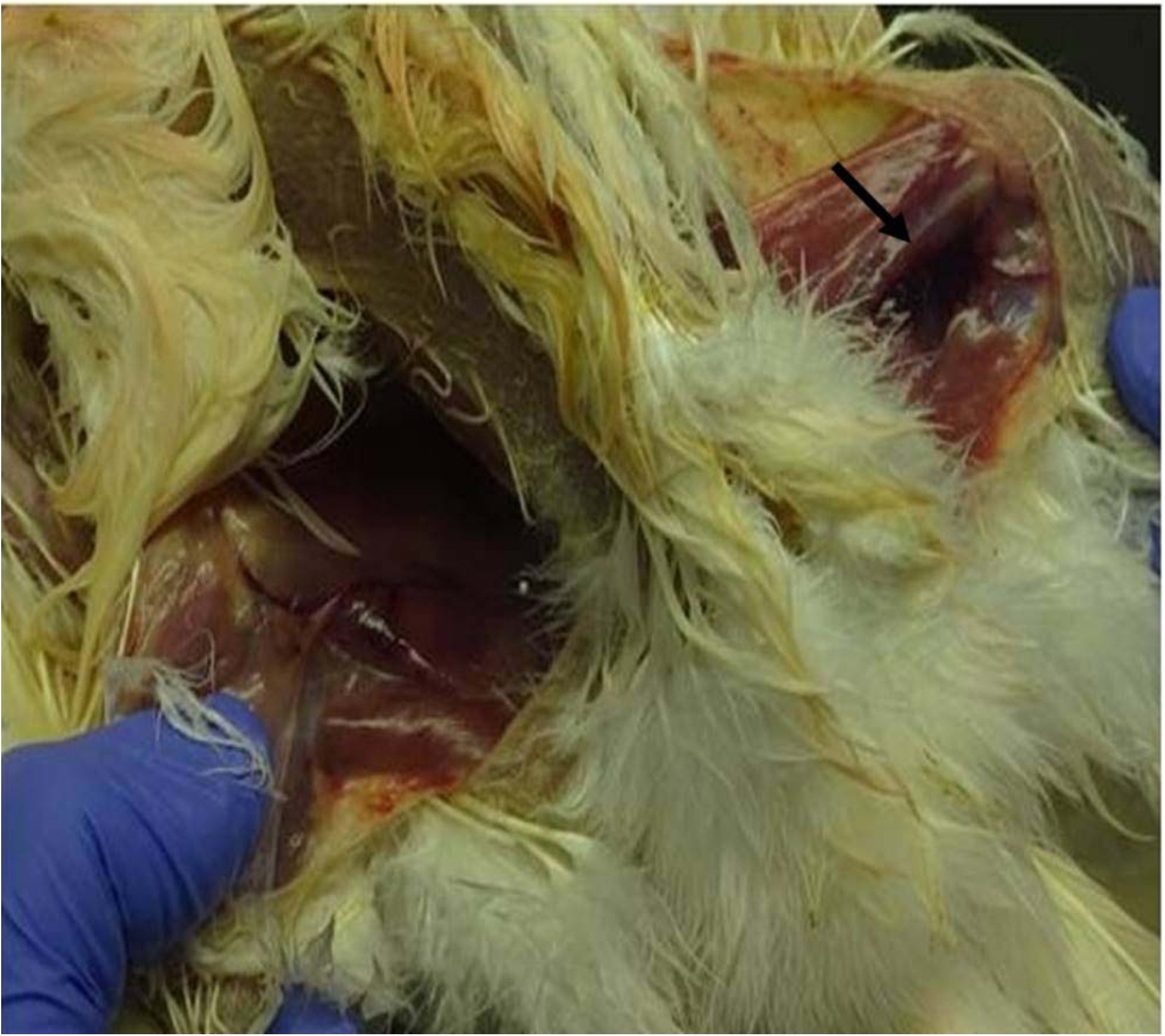
10^6^ EID_50_ group - dead bird (47 hpi) - Thigh muscle: Haemorrhage in thigh muscle (arrow) of dead bird.

Significant histological alterations with varying grades of severity were recorded in various organs of birds infected with 10^4^ and 10^6^ EID_50_ doses of virus at different hpi and in found dead birds (Supplementary Table. 2 & Figs. 7-11). However, no microscopic lesions were observed in birds infected with 10^2^ EID_50_ virus dose and mock infected birds. The predominant changes in nasal cavity were congestion, haemorrhage, heterophil and mononuclear cell infiltration in nasal mucosa and propria - submucosa. Shortening and loss of cilia and mucous secretory cell hypertrophy and hyperplasaia were the major findings in trachea.

**Fig. 7:**
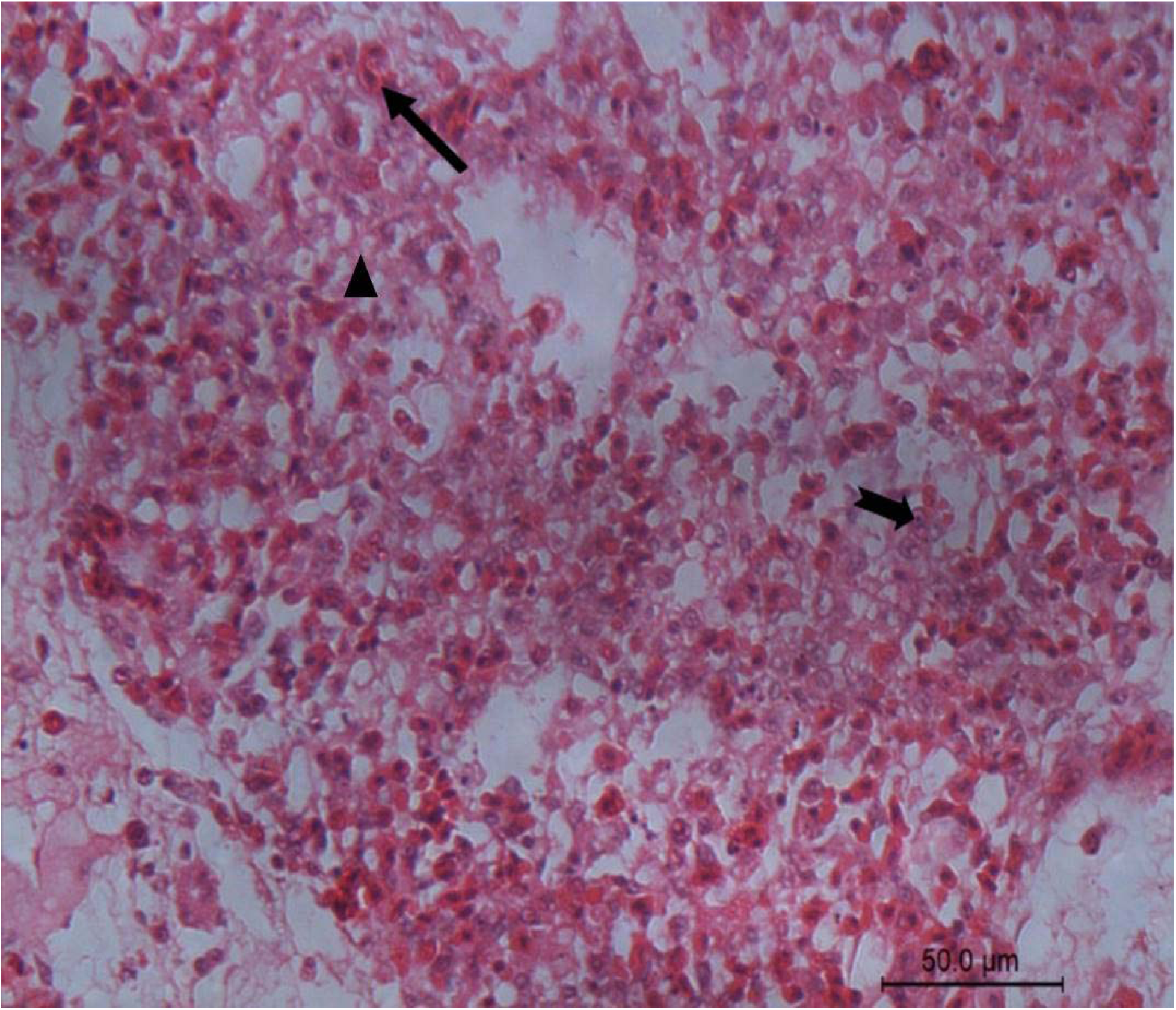
10^6^ EID_50_ group - dead bird (47 hpi) - Lungs: Severe pulmonary consolidation characterized by predominant heterophils infiltration (arrow) along with mononuclear cell proliferation (notched arrow), haemorrahe, and serofibrinous exudates (arrow head) in air capillaries (50 μm).

**Fig. 8:**
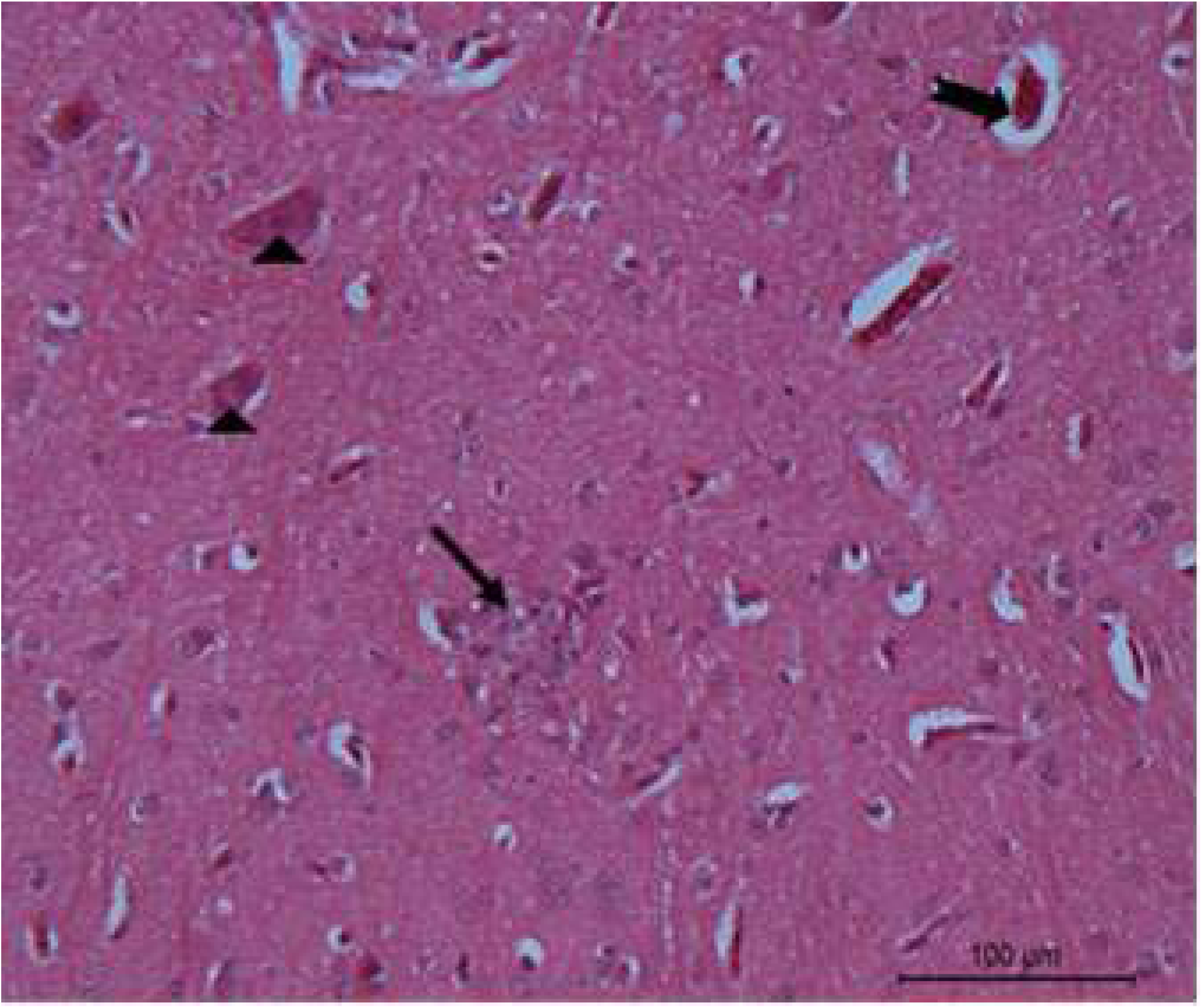
10^6^ EID_50_ group - dead bird (47 hpi) - Brain: Cerebrum showing focal glial nodule (arrow), necrotic or neuronal degeneration (arrow head), and perivascular oedema (notched arrow) (100 μm).

**Fig. 9:**
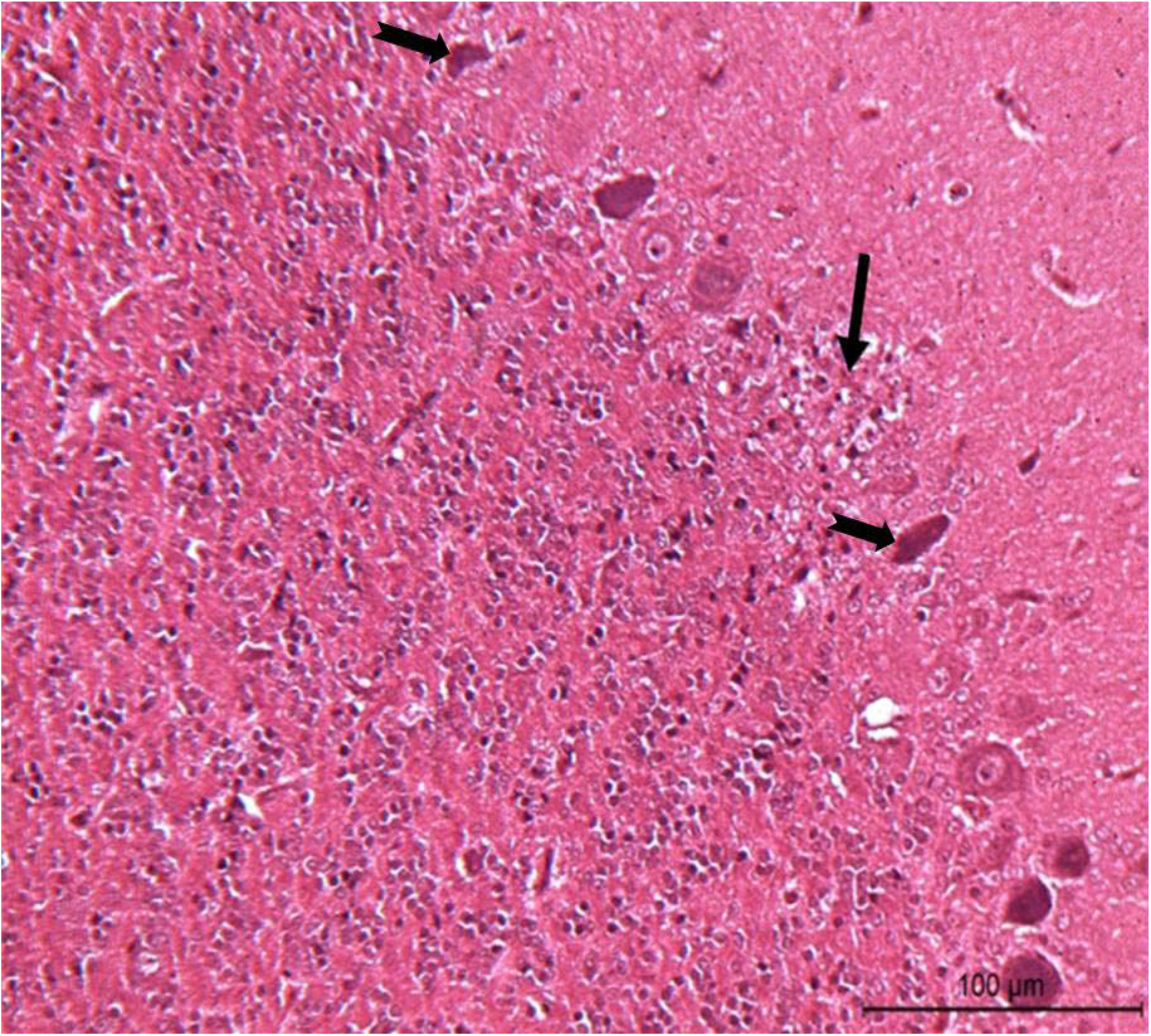
10^6^ EID_50_ group - dead bird (47 hpi) - Brain: Focal malacia (arrow) at the juncture of cerebellar molecular layer and purkinje cell layer along with degenerated (notched arrow) and necrotic (notched arrow) Purkinje neurons (100 μm).

**Fig. 10:**
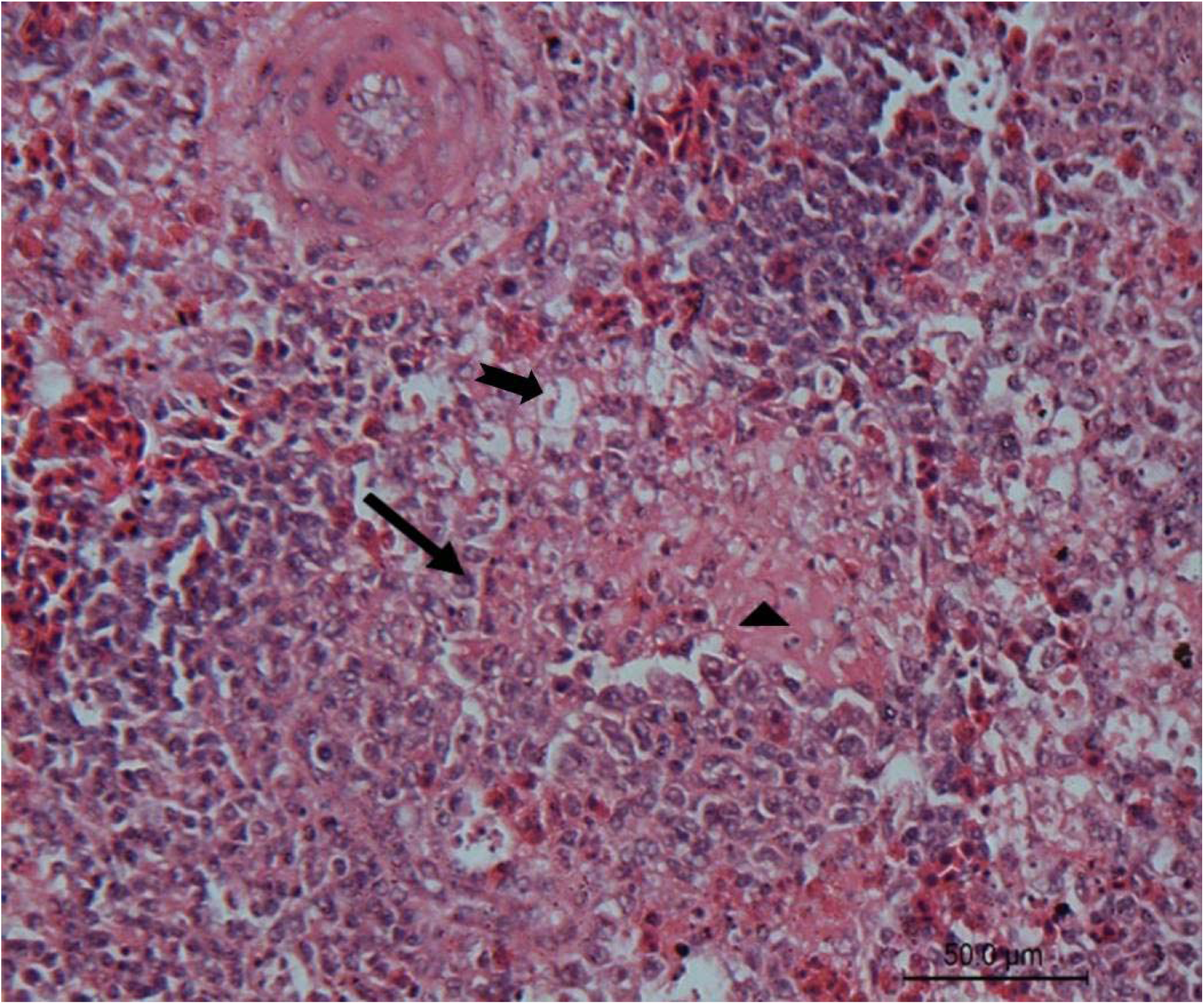
10^6^ EID_50_ group - dead bird (47 hpi) - Spleen: White pulp showing severe depletion of lymphocytes characterized by pinkish necrotic area (arrow head) with pyknotic nuclei and irregular empty spaces of lymphocytolysis (notched arrow) along with phagocytic cell proliferation (arrow) (50 μm).

**Fig. 11:**
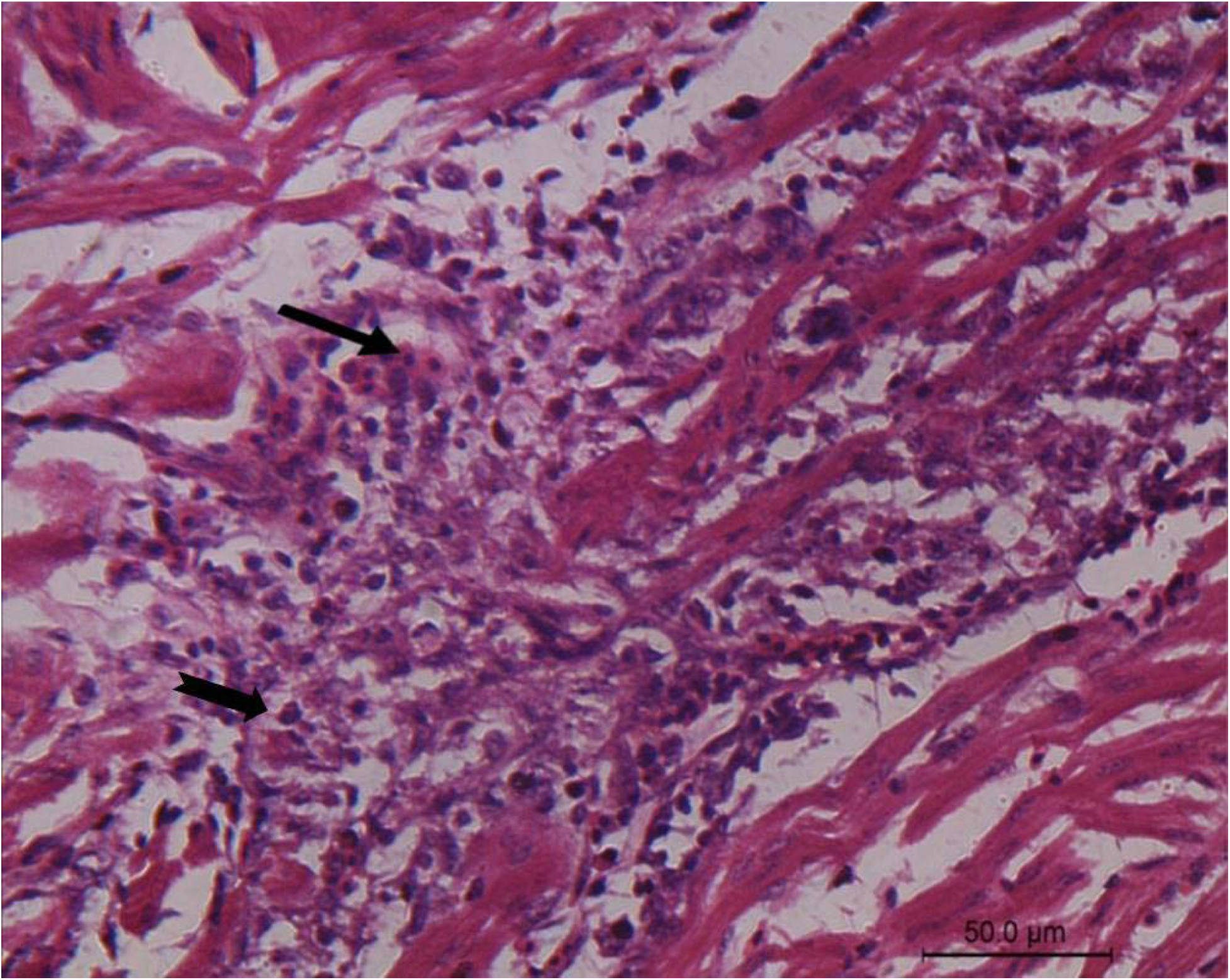
10^4^ EID_50_ group - dead bird (132 hpi) - Heart: Focal necrosis of cardiac myocytes along with infiltration of heterophils (arrow) and mononuclear cells (notched arrow) (50 μm).

Congestion, haemorrhage, serofibrinous exudation, heterophils and mononuclear cell infiltration in air capillaries, bronchi, and peribronchial area were the significant findings in lungs (Fig. 7). In brain (Figs. 8 & 9), perivascular edema, degenerative and necrotic neurons, gliosis, and malacic foci were the significant findings along with paucity of purkinje neurons in cerebellum (Fig. 9). Congestion, hemorrhage, focal hyalinization, necrosis, and heterophil and mononuclear cell infiltration were significant findings in heart (Fig. 11). Sinusoidal dilatation, congestion, haemorrhage, hepatocyte degeneration, and mononuclear cell infiltration were the major findings in liver. Notable findings in kidneys were deneudation and degeneration of Proximal Convoluted Tubules in addition to the congestion.

In intestine, lymphoid depletion of gut lymphoid tissues, necrosis of mucosal and submucosal glands and muscularis layer, and heterophil and mononuclear cell infiltration in mucosa, submucosa and muscularis mucosa were the significant findings. Vacuolar degeneration and necrosis of acinar cells were observed in pancreas. Congestion, lymphoid depletion, and phagocytic cell proliferation were the major findings in spleen (Fig.10). In cortex and medulla of bursa, lymphoid depletion, heterophil infiltration, and phagocytic cell proliferation were observed in addition to the atrophy of follicles. In thymus, congestion of cortical and medullary vessels and lymphoid depletion of cortex depicted the “starry sky appearance”, were the significant findings.

### 3.5. Viral antigen distribution in tissues

Comparison of antigen distribution of all mentioned organs at different hpi from each group was scored and presented (Supplementary Table. 3). No viral antigen was detected in mock infected birds and in 10^2^ EID_50_ group at all the intervals of the study (Supplementay Table 3). In 10^4^ and10^6^ EID_50_ group, viral antigen was demonstrated in various organs except kidney; chondrocytes, endothelial cells, epithelial cells, and mononuclear cells of nasal cavity; chondrocytes and epithelial cells of trachea; endothelial cells and mononuclear cells of lungs (Fig. 12); mononuclear cells of spleen (Fig. 15) thymus, bursa, and intestine; glial cells neurons, and ependymal of cerebrum (Fig. 13); glial cells and granular cells of cerebellum (Fig. 14); cardiac myocytes of heart (Fig. 16); Kupffer cells and hepatocytes of liver; and in acinar cells of pancreas.

**Fig.12:**
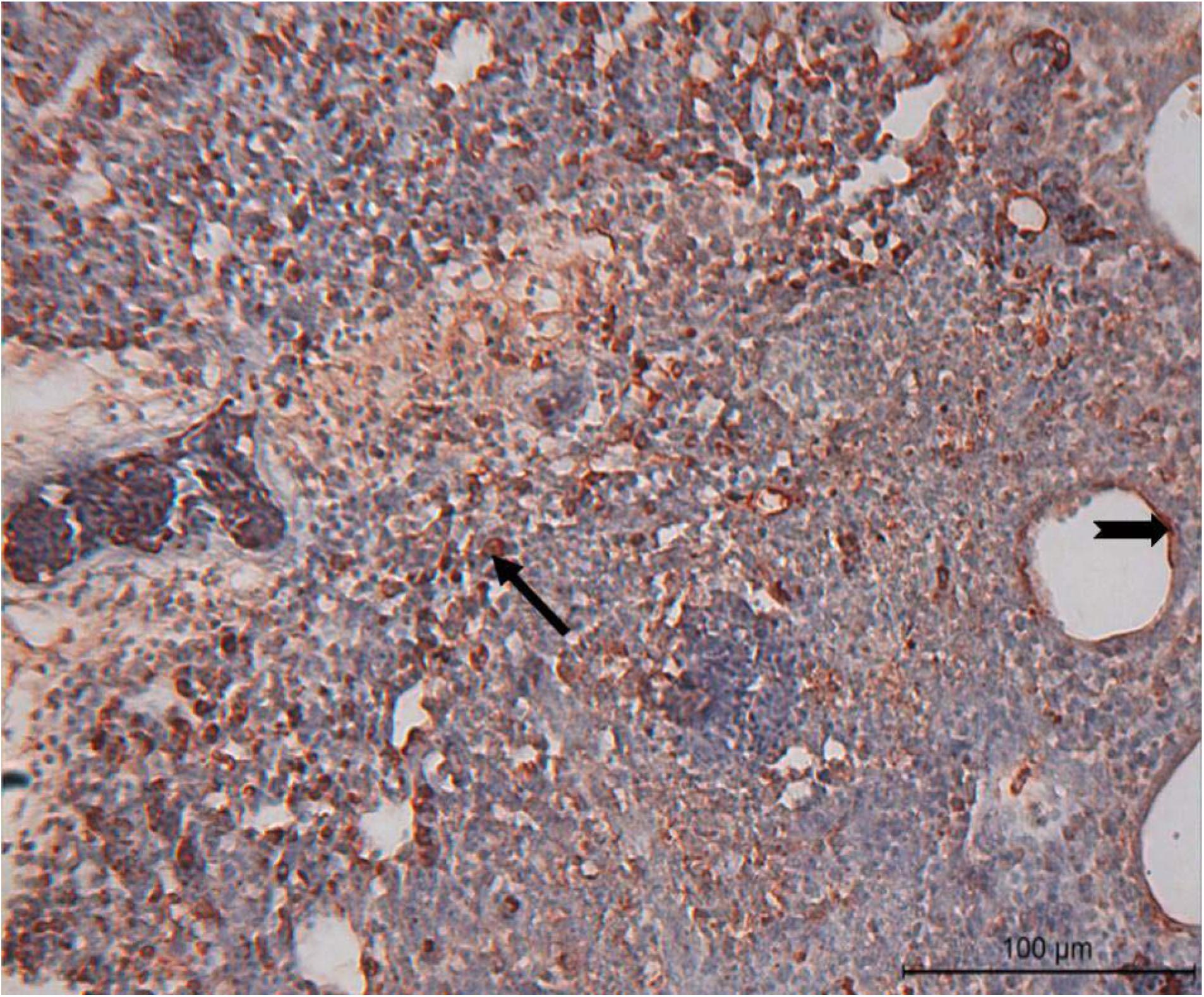
104 EID_50_ group - dead bird (132 hpi) - Lungs: Diffuse viral antigen distribution in monocuclear cells (arrow) of peribronchiolar and aircapillary area and in endothelium (notched arrow) (100 μm).

**Fig. 13:**
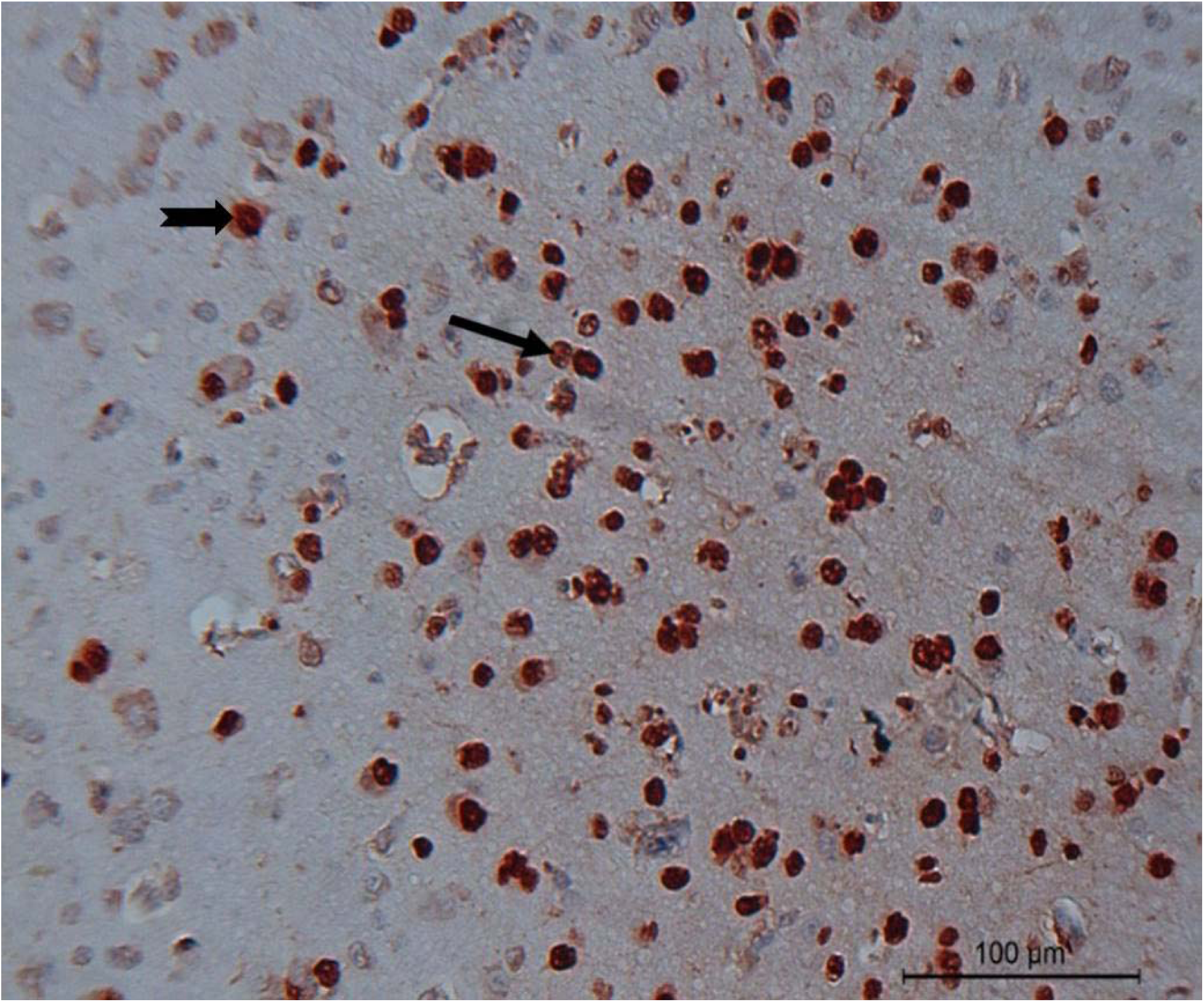
10^4^ EID_50_ group - dead bird (132 hpi) - Brain: Cerebrum showing positive staining of viral antigen in glial cells (arrow) and neurons (notched arrow) (100 μm).

**Fig. 14:**
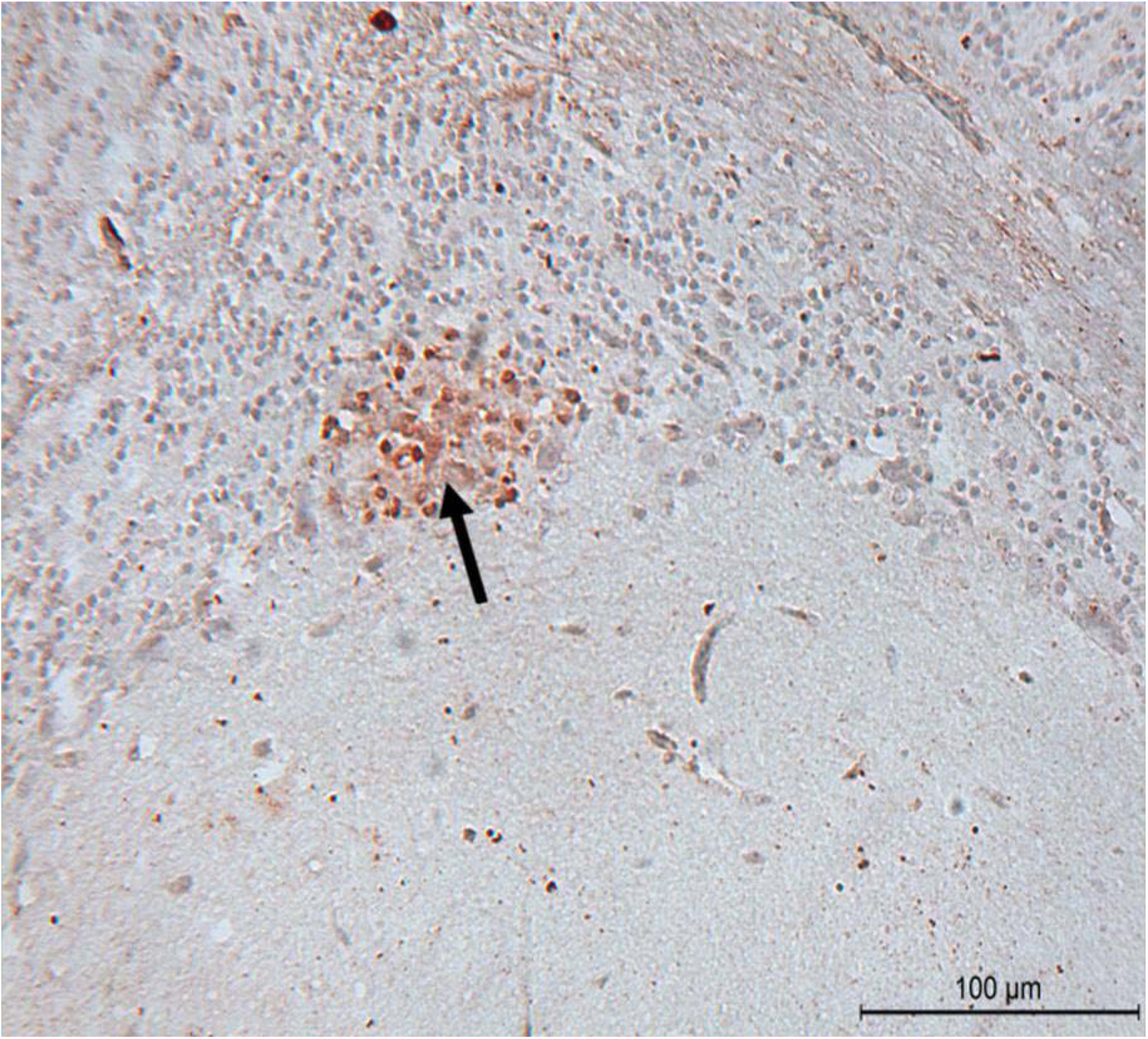
10^6^ EID_50_ group - dead bird (47 hpi) - Brain: Cerebellum showing positive viral antigen signal of malacic foci (arrow) at the juncture of purkinje cell layer and granular cell layer (100 μm).

**Fig. 15:**
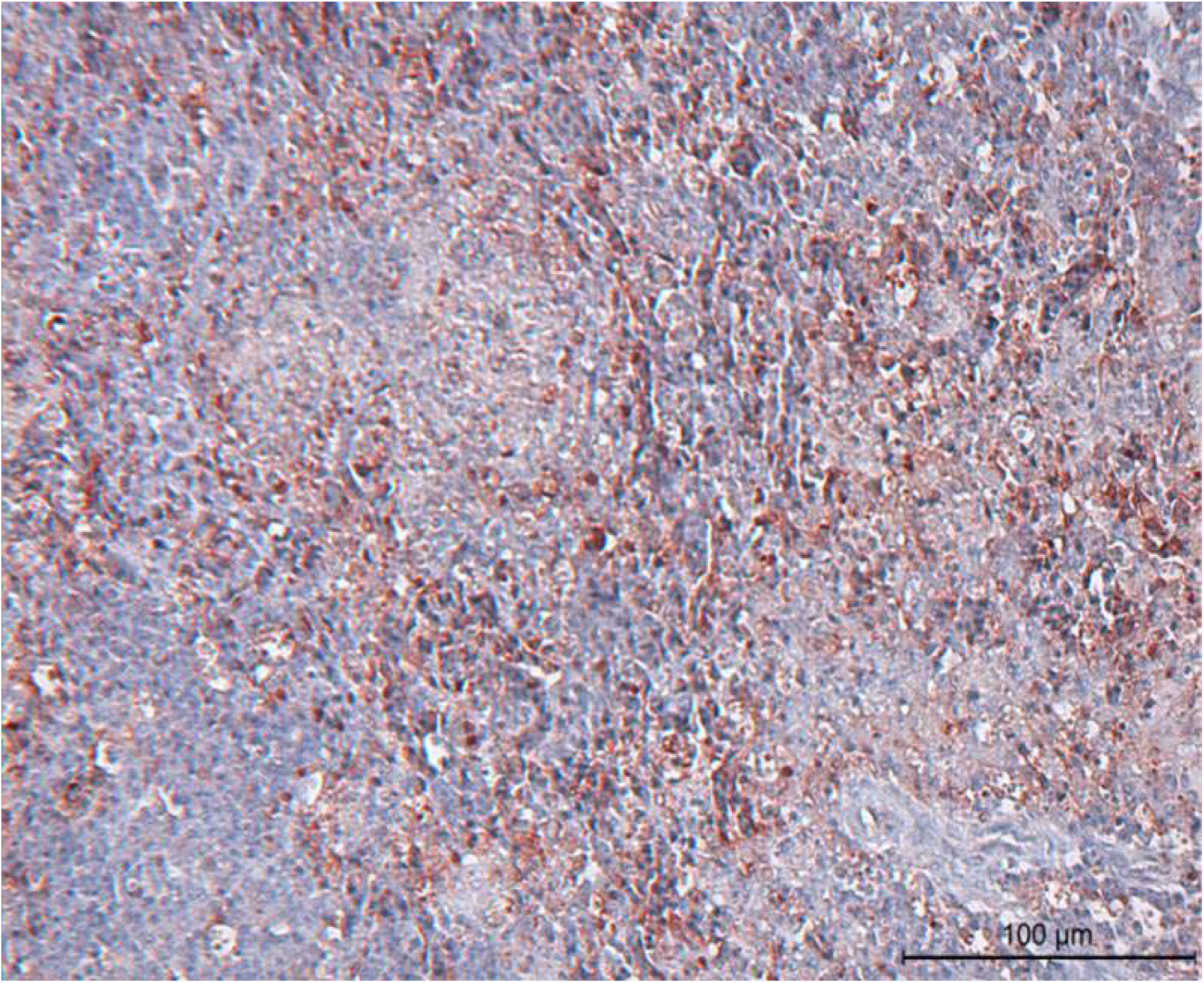
10^6^ EID_50_ group - dead bird (47 hpi) - Spleen: Diffuse distribution of viral antigen in mononuclear cells (100 μm).

**Fig. 16:**
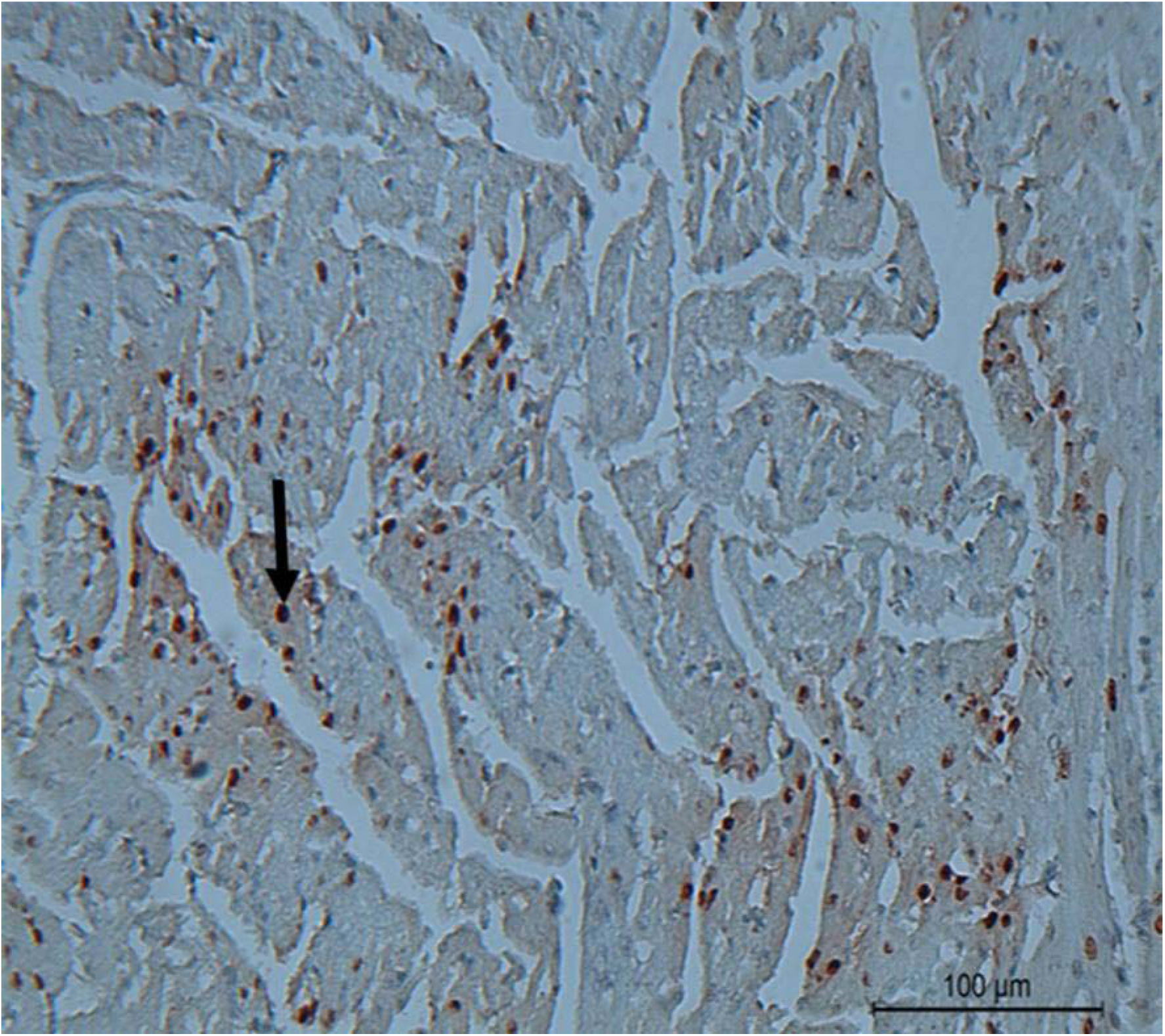
10^6^ EID_50_ group - dead bird (47 hpi) - Heart: Showing strong diffuse / multi focal viral antigen positive signals in myocardium (arrow) (100 μm).

## 4. Discussion

Global presence of HPAI H5 virus constitute threat to animal and potentially to human health. H5Nx Gs/GD lineage clade, 2.3.4.4 viruses have caused huge mortality of domestic and wild birds and economic loss in affected countries. Few studies have showed that these viruses pose low zoonotic potential [37–39]. In india, outbreaks of HPAI, H5N8 virus were found in wild birds, chicken and domestic ducks. In the present study, virus of chicken origin, H5N8 virus A/chicken/India/11CA01/2016, was used to investigate the pathobiology of H5N8 virus in chicken model.

Differences in progression of disease and mortality pattern were observed between10^4^ and 10^6^ EID_50_ groups after inoculation of virus. Duration of the infection, the organ or organ systems affected, and the duration of infection were the factors associated with clinical signs in HPAI [40]. Dullness, huddling, anorexia, prostration, dropping of eyelids and the nervous signs in the present study were consistent with previous description [28–30, 40]. Delayed death of birds in 10^4^ EID_50_ group attribute more pronounced nervous signs, such as torticollis and circling than birds found dead with the 10^6^ EID_50_ dose.

HPAI viruses caused high mortality rate in chicken as well as in other gallinaceous birds but the duration is shorter in chicken and vary with individual HPAI strain [40]. MDT of previously reported indian isolate of HPAIV H5N1 with dose 10^2^ EID_50_ was 4.57 days [41]. So, in current study absence of clinical signs and mortality throughout the course and absence of seroconversion at 14 dpi sacrificed birds in 10^2^ EID_50_ group indicate insufficiency of dose inocula to initiate and induce the systemic infection in birds. It is evident that individual variation in the HPAI virus subtype to cause disease in particular host species with same dose. Results are consistent with previous reports of H5N8 with the dose, 10^2^ EID_50_ [28, 30] in chicken. Earlier reports of MDT of indian HPAI, H5N1 isolates with dose of 10^4^ EID_50_ was 3.6 days [41], is comparable to the present study result, 3.2 days. Park *et al.* (2021) [25] reported MDT of 7.4 days in the experimental infection of chickens with the inocula 10^4^ EID_50_. On other hand, experimental challenge of HPAI, H5N8 virus with 10^4^ EID_50_ dose inocula did not cause any mortality in chicken [30] attributed to variation in the virulence of different strains of HPAI virus subtype for the same host species. Reports of MDT of 10^6^ EID_50_ dose inocula showed more than 3 (3.2-4.8) days duration in experimentally challenged chickens [24, 27, 28]. However, varying duration of MDT in chickens infected by 10^6-6.5^ EID_50_ of H5N8 virus were reported with MDT, 2-2.2 [30], 2.5 [26], and 4.3 days[25], which was comparable to the present study result, 2 days. Thus, our results showed highly virulent nature of indian HPAI, H5N8 isolates in chicken. Earlier onset of mortality, higher cumulative percent mortality (100%), and lesser MDT of 10^6^ EID_50_ group than dose group, 10^4^ EID_50_ denotes dose dependent outcome of the clinical disease and variation in the progression of disease.

CID_50_ under experimental conditions is a host and virus dependent factor, which is a predictor of the infectivity and adaption to specific host [22]. In our study, the resultant CID_50_ of A/chicken/India/11CA01/2016 (H5N8) virus was 10^4.5^ EID_50_. This findings are comparable to the result of 10^4.4^ [28] and 10^5^ [30] EID_50_, respectively. HPAI virus, AsianA/HongKong/486/1997(H5N1) is known to be one of the most virulent and poultry adapted HPAI viruses [42, 43] having CID_50_ of 10^2.4^ EID_50_ [22]. Moreover, CID_50_ of 10^4.7^ EID_50_ was proposed as upper cut off for the influenza viruses to initiate infection and set the conditions for virus strain adaption and transmission for the specific host species in field conditions [22]. With reference to this, in current study, results indicate that this novel HPAIV, H5N8 viruses showing poor adaption to chicken [28, 30].

Replication of HPAI virus with high titre in respiratory and intestinal tracts of chickens cause shedding of virus in nasal secretions and faeces [40]. In reference to virus replication and shedding of HPAI virus, amount of virus in excretion has a direct impact on the degree of environnmental contamination and subsequent bird-to-bird transmission and final farm-to-farm spread [28]. In 10^2^ EID_50_ dose group, absence of shedding from 6 hpi to 14 dpi correlated with insufficiency of the dose inocula to replicate in the intestinal tract and respiratory system. These findings were corroborated by the results of the previous study with 10^2^ EID_50_ of H5N8 virus in chicken [28, 30]. High viral titres of oropharyngeal and cloacal shedding both in 10^4^ and 10^6^ EID_50_ groups at all dpi were consistent with previous reports of experimental studies of H5N8 infection in chicken [25, 28, 30]. Absence of viral sheddings in survived birds after 5 dpi in 10^4^ EID_50_ group indicates lack of systemic replication of virus to initiate infection, which is similar to that of 10^2^ EID_50_ group.

The dissemination of HPAI viruses generally progresses from an initial respiratory infection to the circulatory system and subsequently other systems in a sequential and time dependent manner [44]. Initial detection of virus in nasal cavity and trachea in 10^2^ EID_50_ group denotes the local replication of virus in birds. Further, failure of 10^2^ EID_50_ dose to reach after 6 hpi till 14 dpi in all organs indicates insufficient dose inocula to reach and replicate systemically. Absence of viral RNA in various organs with 10^2^ EID_50_ dose was concordant with earlier findings [28, 30]. High viral load in multiple organs such as nasal cavity, trachea, lungs, brain, spleen, heart, liver, kidney, intestine, pancreas were noticed with the dose inocula 10^4^ and 10^6^ EID_50_ in the present study. The quantity of high virus copy numbers in various organ is directly associated with death of the individual birds. The viral copy numbers of various organs varied with high (10^5^-10^10^/g tissue) to very high (> 10^10^ / g tissue) titres at different hpi. Highest viral copy numbers in 10^4^ EID_50_ were noticed in nasal cavity, kidney, spleen, and heart at 6 & 12, 24, 48, and 53-232 hpi, respectively. In 10^6^ EID_50_ group, highest viral RNA copy numbers were noticed in nasal cavity, lungs, and heart at 6 & 12, 24 & 48 (40-47) hpi, respectively. The results were in concurrence with previous reports which had reported varying high viral titres in multiple organs of chicken experimentally infected with HPAI, H5N8 viruses. [25,28,30]. This variation in viral copy numbers is due to the binding properties of influenza virus HA to glycan receptors, which in turn affects organ tropism and virulence of virus in the host species [45]. Thus, our results indicate the systemic replication of HPAI, H5N8 virus in affected chicken. Further, survived birds in both 10^2^ and 10^4^ EID_50_ group, might have been cleared the virus early by local immune mechanism from upper respiratory passage before reaching the systemic path. It is correlated to no detectable viral RNA copies in oropharyngeal and cloacal shedding and in multiple organs subjected for virus detection, and absence of seroconversion at termination, 14 dpi.

Severity of lesions in various organs was directly related with viral replication, which ultimately causes mortality in affected chickens. Gross lesions such as congestion, haemorrhage, and odema in the various above-mentioned organs were consistent with earlier reports in addition to the finding of pale liver and whitish foci of spleen [28,30,40,41]. Microscopic lesions in chickens are more consistent than gross lesions in HPAI cases [40]. Microscopically lesions were of congestion, haemorrhage and inflammatory changes in nasal cavity and lungs, which were consistent with earlier findings of experimental infection with HPAI viruses [28, 30, 40–42]. Loss of cilia and mucous secretory cell hypertrophy and hyperplasia in trachea were comparable with earlier reports [41]. Degenerative lesions, vascular changes, and necrotic and inflammatory findings of liver and heart were similar to described by [28, 30, 40, 41]. Deneudation of Proximal Convoluted Tubules in cortex along with vascular and degenerative changes in kidney were consistent findings with earlier reports [41]. Necrotic and inflammatory lesions and lymphoid deletion of lymphoid aggregates of intestine were similar that of earlier findings [40, 42]. Vacuolar degeneration and acinar cell necrosis of pancreas were consistent with earlier descriptions [40]. Lymhoid depletion and phagocytic cell proliferation of spleen and bursa and heterophil infiltration and atrophy of bursal follicles were similar to those described for infection of chicken with HPAIV infections [40–42]. Vascular changes and lymphoid deletion of thymus were coincided with previous study reports [40–42].

Viral antigen detection in various cells of multiple organs coincides with varying degree of severity of microscopic lesions in 10^4^ and 10^6^ EID_50_ group. Moreover, staining frequency of tissues with viral antign was varied and scored at different hpi. Similar findings of widespread staining of HPAI viral antigen in multiple organs was described earlier [28, 30, 40, 42]. Widespread endothelial cell staining in multiple organs was not observed in the present study [28, 46] and such staining was restricted to nasal cavity and lungs only. Activation of coagulation cascade in HPAI virus infected birds by replication in endothelial cells and/or monocytes/ macrophage induces Disseminated Intravascular Coagulation (DIC) [47]. Thus, our results indicates that absence of widespread endothelial staining in multiple organs lacks the DIC which attributes variation in replication and pathogenesis of Gs/Gd lineage clade, 2.3.4.4b HPAI, H5N8 virus from other HPAI, H5N1 virus.

Thus, the study concludes that Indian HPAI, H5N8 isolates of clade 2.3.4.4b are highly pathogenic in nature to chicken by affecting most organs systemically. CID_50_ of this H5N8 virus indicates poor adaption in chicken and it implies poor transmission possibility of this virus for host species in field condition. Pathological features were similar as that of HAPIV, H5N1 with the exception of Disseminated Intravascular Coagulation at the dose level 10^4^ and 10^6^ EID_50_ and lack of endothelial staining except nasal cavity and lungs implies variation in the pathogenesis of this novel virus from HPAI, H5N1 virus. Hence, further concern is required to elucidate the pathobiology of these viruses in various bird species.

## Supporting information

https://docs.google.com/document/d/1MYaACePdfCa1YULfdp-2AmA7SID28Erv/edit?usp=drive_link&ouid=105182866489515572078&rtpof=true&sd=true

## CRediT author statement

Boopathi Ponnusamy: Conceptualization, Methodology, Data curation, Writing - original draft. Manoj Kumar: Conceptualization, Methodology, Supervision. Harshad Vinayakrao Murugkar: Data curation, Writing - original draft. Shanmugasundaram Nagarajan: Data curation, Writing - original draft, Writing - review & editing. Chakradhar Tosh: Visualization, Investigation. Sivasankar Panickan: Data curation, Writing - original draft, Writing - review & editing. Dhruv Desai: Data curation, Writing - original draft, Writing - review & editing. Semmannan Kalaiyarasu: Conceptualization, Methodology. Dhanapal Senthil Kumar: Data curation, Writing - original draft. Siddharth Gautam: Supervision, Writing - review & editing. Vijendra Pal Singh: Supervision, Writing - review & editing. Aniket Sanyal: Writing - review & editing.

## Acknowledgements

We would like to acknowledge the grant from Indian Council of Agricultural Research (ICAR). We also thank the Directors of ICAR-Indian Veterinary Research Institute (ICAR-IVRI), Bareilly and ICAR- National Institute of High Security Animal Diseases (ICAR-NIHSAD), Bhopal for providing necessary facilities to carry out this work.

## Supplementary documents

**Supplementary Table. 1:**
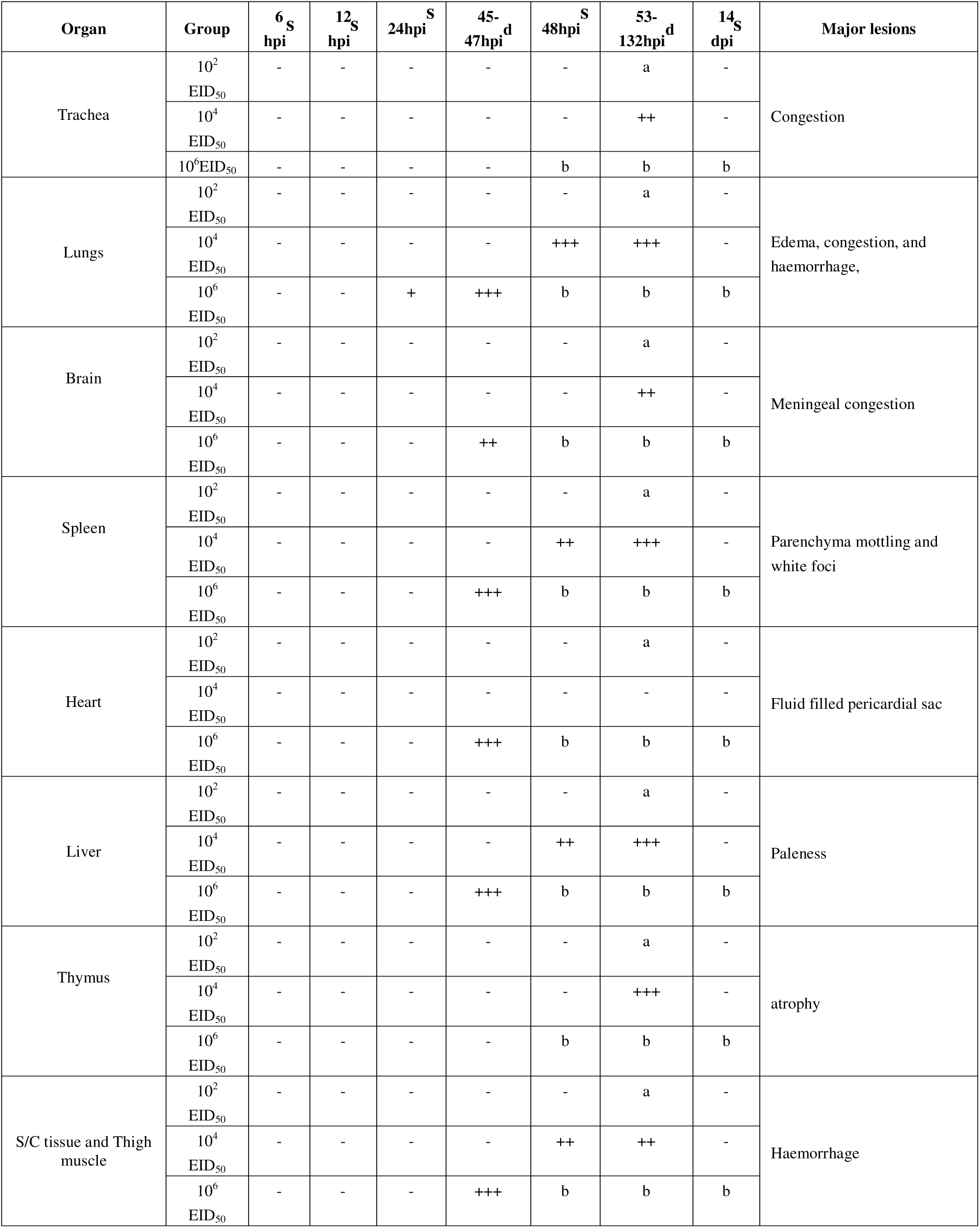
Gross lesions* at different intervals in chicken inoculated with A/chicken/India/n11CA01/2016 (H5N8) virus with 10^2^, 10^4^ and 10^6^ EID50 * - = None, ± = Minimal, + = Mild, ++ = Moderate, and +++ = Severe; a - No birds were euthanized or died at the respective interval; b - All birds died before 48 hpi in 10^6^ EID50 group; s - Birds were euthanized at the respective interval; d - Dead birds.

**Supplementary Table. 2:**
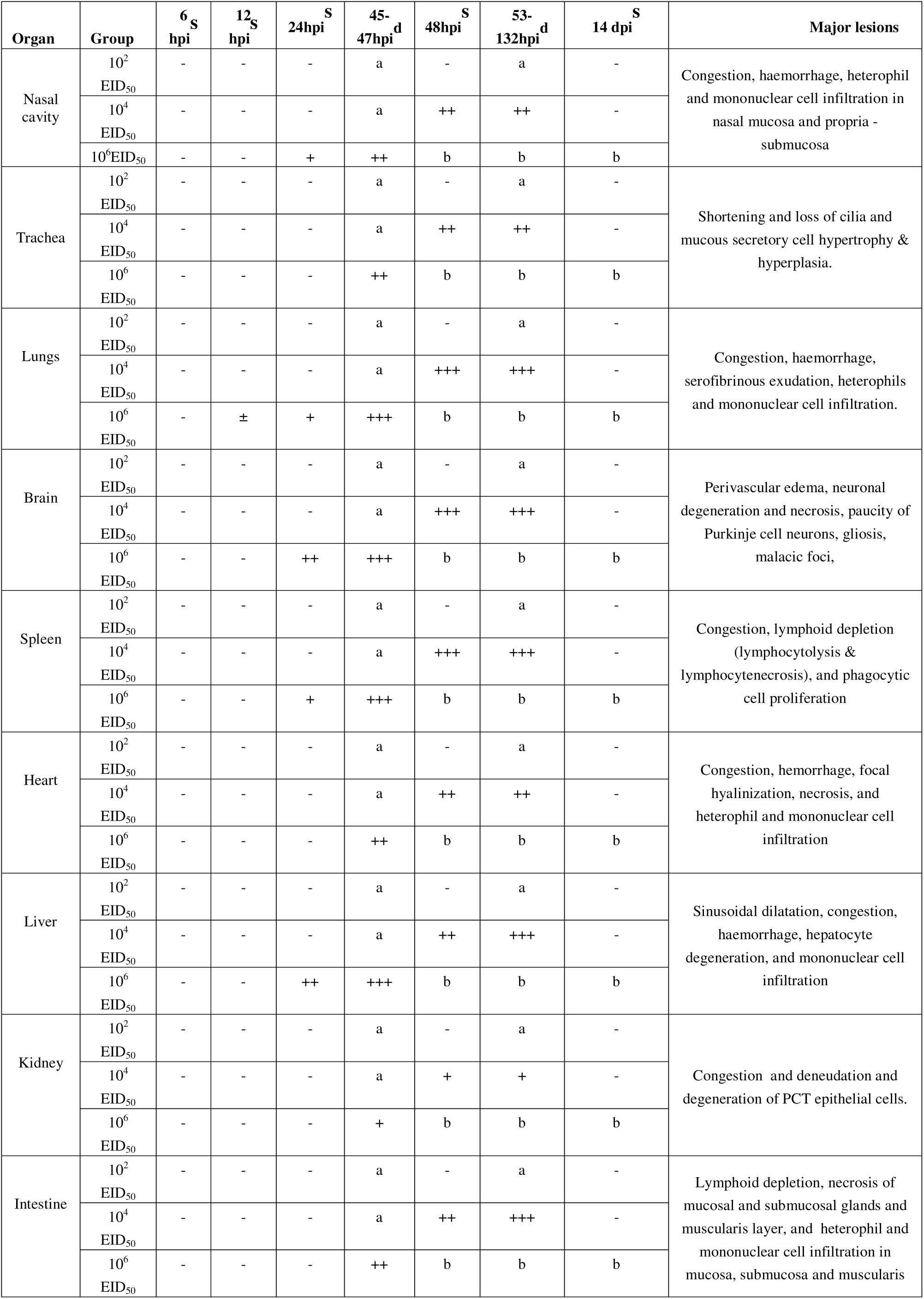
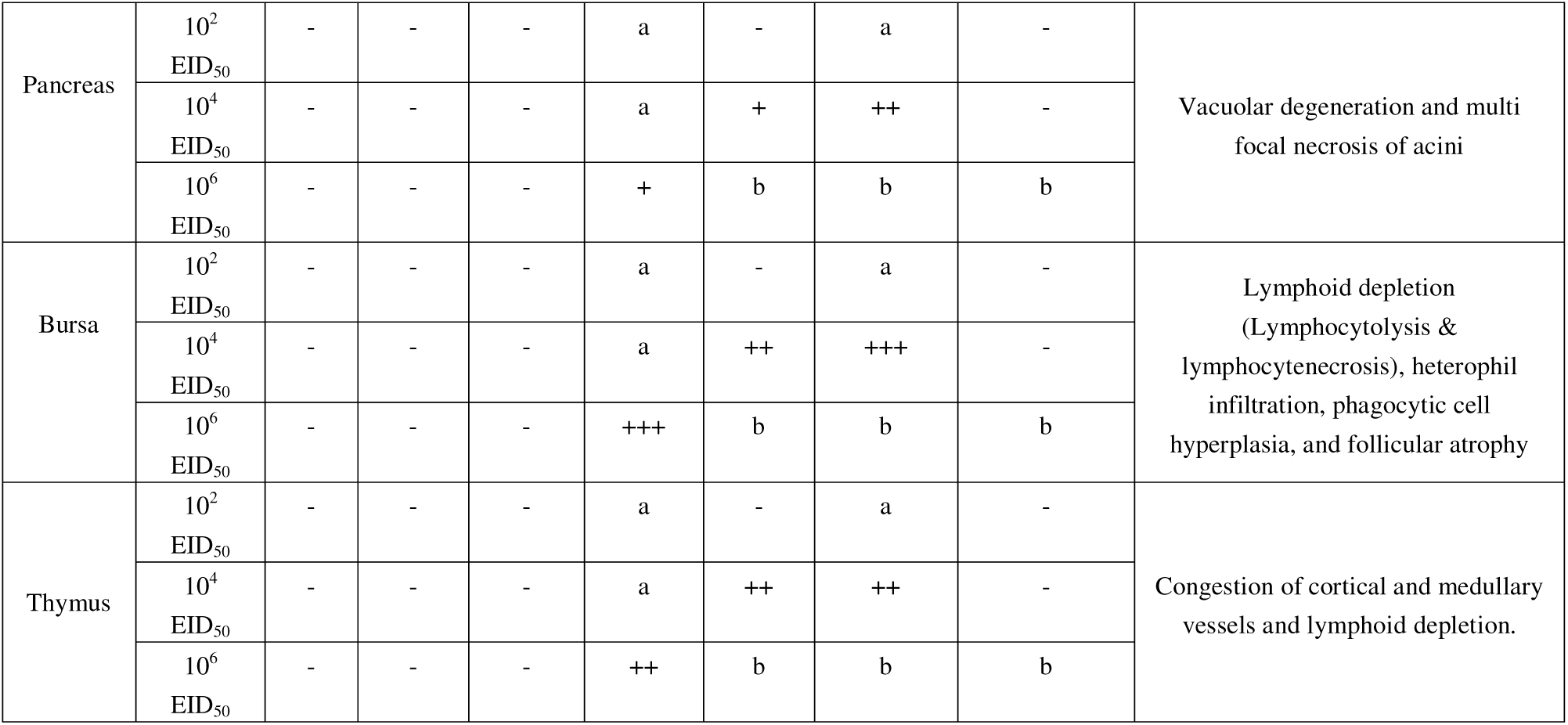
Histopathological lesions* at different intervals in chicken inoculated with A/chicken/India/n11CA01/2016 (H5N8) virus with 10^2^, 10^4^ and 106 EID^50^. * - = None, ± = Minimal, + = Mild, ++ = Moderate, and +++ = Severe; a - No birds were euthanized or died at the respective interval; b - All birds died before 48 hpi in 106 EID50 group; s - Birds were euthanized at the respective interval; d - Dead birds

**Supplementary Table. 3:**
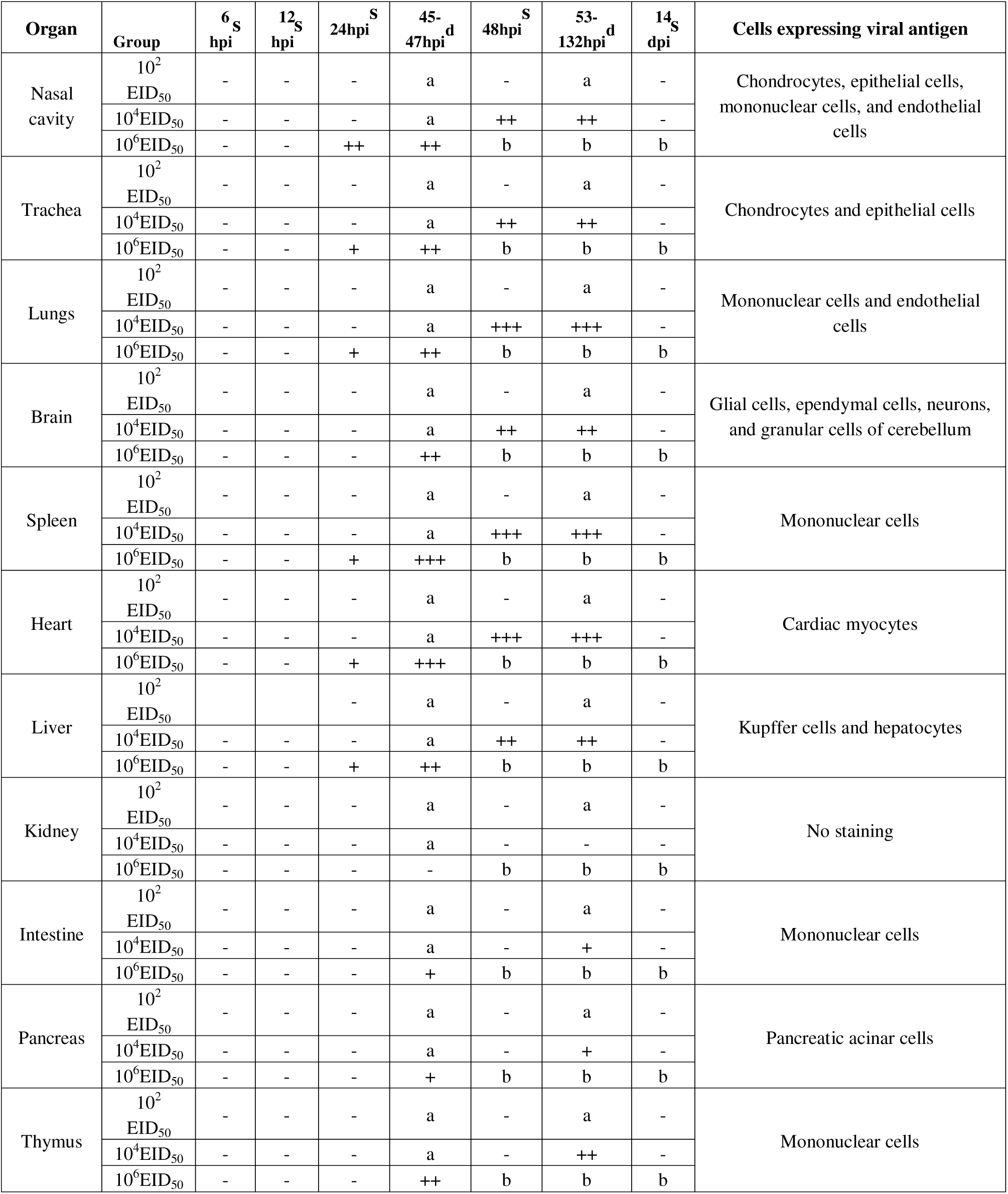
Distribution* of H5N8 antigen in virus infected tissues of chicken with different doses, 10^2^, 10^4^ and 10^6^ EID_50_ (Immunohistochemistry). * - = None; + = Infrequent staining, ++ = Frequent staining, and +++ = Diffuse staining; a - No birds were euthanized or died; b - All birds died before 48 hpi in 10^6^ EID50 group; s - Birds were euthanized at the respective interval; d - Dead birds.

