## Supplementary material for "Pathology of dose dependent inocula of H5N8 avian influenza virus in experimentally infected chicken": https://docs.google.com/document/d/1MYaACePdfCa1YULfdp-2AmA7SID28Erv/edit?usp=drive_link&ouid=105182866489515572078&rtpof=true&sd=true

| **Organ** | **Group** | **6 hpi^s^** | **12 hpi^s^** | **24hpi^s^** | **45-47hpi^d^** | **48hpi^s^** | **53-132hpi^d^** | **14 dpi^s^** | **Major lesions** |
| --- | --- | --- | --- | --- | --- | --- | --- | --- | --- |
| Trachea | 10^2^ EID_50_ | - | - | - | - | - | a | - | Congestion |
|  | 10^4^ EID_50_ | - | - | - | - | - | ++ | - |  |
|  | 10^6^EID_50_ | - | - | - | - | b | b | b |  |
| Lungs | 10^2^ EID_50_ | - | - | - | - | - | a | - | Edema, congestion, and haemorrhage, |
|  | 10^4^ EID_50_ | - | - | - | - | +++ | +++ | - |  |
|  | 10^6^ EID_50_ | - | - | + | +++ | b | b | b |  |
| Brain | 10^2^ EID_50_ | - | - | - | - | - | a | - | Meningeal congestion |
|  | 10^4^ EID_50_ | - | - | - | - | - | ++ | - |  |
|  | 10^6^ EID_50_ | - | - | - | ++ | b | b | b |  |
| Spleen | 10^2^ EID_50_ | - | - | - | - | - | a | - | Parenchyma mottling and white foci |
|  | 10^4^ EID_50_ | - | - | - | - | ++ | +++ | - |  |
|  | 10^6^ EID_50_ | - | - | - | +++ | b | b | b |  |
| Heart | 10^2^ EID_50_ | - | - | - | - | - | a | - | Fluid filled pericardial sac |
|  | 10^4^ EID_50_ | - | - | - | - | - | - | - |  |
|  | 10^6^ EID_50_ | - | - | - | +++ | b | b | b |  |
| Liver | 10^2^ EID_50_ | - | - | - | - | - | a | - | Paleness |
|  | 10^4^ EID_50_ | - | - | - | - | ++ | +++ | - |  |
|  | 10^6^ EID_50_ | - | - | - | +++ | b | b | b |  |
| Thymus | 10^2^ EID_50_ | - | - | - | - | - | a | - | atrophy |
|  | 10^4^ EID_50_ | - | - | - | - | - | +++ | - |  |
|  | 10^6^ EID_50_ | - | - | - | - | b | b | b |  |
| S/C tissue and Thigh muscle | 10^2^ EID_50_ | - | - | - | - | - | a | - | Haemorrhage |
|  | 10^4^ EID_50_ | - | - | - | - | ++ | ++ | - |  |
|  | 10^6^ EID_50_ | - | - | - | +++ | b | b | b |  |

* - = None, ± = Minimal, + = Mild, ++ = Moderate, and +++ = Severe; a - No birds were euthanized or died at the respective interval; b - All birds died before 48 hpi in 10^6^ EID_50_ group; s - Birds were euthanized at the respective interval; d - Dead birds

| **Organ** | **Group** | **6 hpi^s^** | **12 hpi^s^** | **24hpi^s^** | **45-47hpi^d^** | **48hpi^s^** | **53-132hpi^d^** | **14 dpi^s^** | **Major lesions** |
| --- | --- | --- | --- | --- | --- | --- | --- | --- | --- |
| Nasal cavity | 10^2^ EID_50_ | - | - | - | a | - | a | - | Congestion, haemorrhage, heterophil and mononuclear cell infiltration in nasal mucosa and propria - submucosa |
|  | 10^4^ EID_50_ | - | - | - | a | ++ | ++ | - |  |
|  | 10^6^EID_50_ | - | - | + | ++ | b | b | b |  |
| Trachea | 10^2^ EID_50_ | - | - | - | a | - | a | - | Shortening and loss of cilia and mucous secretory cell hypertrophy & hyperplasia. |
|  | 10^4^ EID_50_ | - | - | - | a | ++ | ++ | - |  |
|  | 10^6^ EID_50_ | - | - | - | ++ | b | b | b |  |
| Lungs | 10^2^ EID_50_ | - | - | - | a | - | a | - | Congestion, haemorrhage, serofibrinous exudation, heterophils and mononuclear cell infiltration. |
|  | 10^4^ EID_50_ | - | - | - | a | +++ | +++ | - |  |
|  | 10^6^ EID_50_ | - | ± | + | +++ | b | b | b |  |
| Brain | 10^2^ EID_50_ | - | - | - | a | - | a | - | Perivascular edema, neuronal degeneration and necrosis, paucity of Purkinje cell neurons, gliosis, malacic foci, |
|  | 10^4^ EID_50_ | - | - | - | a | +++ | +++ | - |  |
|  | 10^6^ EID_50_ | - | - | ++ | +++ | b | b | b |  |
| Spleen | 10^2^ EID_50_ | - | - | - | a | - | a | - | Congestion, lymphoid depletion (lymphocytolysis & lymphocytenecrosis), and phagocytic cell proliferation |
|  | 10^4^ EID_50_ | - | - | - | a | +++ | +++ | - |  |
|  | 10^6^ EID_50_ | - | - | + | +++ | b | b | b |  |
| Heart | 10^2^ EID_50_ | - | - | - | a | - | a | - | Congestion, hemorrhage, focal hyalinization, necrosis, and heterophil and mononuclear cell infiltration |
|  | 10^4^ EID_50_ | - | - | - | a | ++ | ++ | - |  |
|  | 10^6^ EID_50_ | - | - | - | ++ | b | b | b |  |
| Liver | 10^2^ EID_50_ | - | - | - | a | - | a | - | Sinusoidal dilatation, congestion, haemorrhage, hepatocyte degeneration, and mononuclear cell infiltration |
|  | 10^4^ EID_50_ | - | - | - | a | ++ | +++ | - |  |
|  | 10^6^ EID_50_ | - | - | ++ | +++ | b | b | b |  |
| Kidney | 10^2^ EID_50_ | - | - | - | a | - | a | - | Congestion and deneudation and degeneration of PCT epithelial cells. |
|  | 10^4^ EID_50_ | - | - | - | a | + | + | - |  |
|  | 10^6^ EID_50_ | - | - | - | + | b | b | b |  |
| Intestine | 10^2^ EID_50_ | - | - | - | a | - | a | - | Lymphoid depletion, necrosis of mucosal and submucosal glands and muscularis layer, and heterophil and mononuclear cell infiltration in mucosa, submucosa and muscularis |
|  | 10^4^ EID_50_ | - | - | - | a | ++ | +++ | - |  |
|  | 10^6^ EID_50_ | - | - | - | ++ | b | b | b |  |
| Pancreas | 10^2^ EID_50_ | - | - | - | a | - | a | - | Vacuolar degeneration and multi focal necrosis of acini |
|  | 10^4^ EID_50_ | - | - | - | a | + | ++ | - |  |
|  | 10^6^ EID_50_ | - | - | - | + | b | b | b |  |
| Bursa | 10^2^ EID_50_ | - | - | - | a | - | a | - | Lymphoid depletion (Lymphocytolysis & lymphocytenecrosis), heterophil infiltration, phagocytic cell hyperplasia, and follicular atrophy |
|  | 10^4^ EID_50_ | - | - | - | a | ++ | +++ | - |  |
|  | 10^6^ EID_50_ | - | - | - | +++ | b | b | b |  |
| Thymus | 10^2^ EID_50_ | - | - | - | a | - | a | - | Congestion of cortical and medullary vessels and lymphoid depletion. |
|  | 10^4^ EID_50_ | - | - | - | a | ++ | ++ | - |  |
|  | 10^6^ EID_50_ | - | - | - | ++ | b | b | b |  |

* - = None, ± = Minimal, + = Mild, ++ = Moderate, and +++ = Severe; a - No birds were euthanized or died at the respective interval; b - All birds died before 48 hpi in 10^6^ EID_50_ group; s - Birds were euthanized at the respective interval; d - Dead birds

Table 3. Distribution * of H5N8 antigen in virus infected tissues of chicken with different doses, 10^2^ , 10^4^ and 10^6^ EID_50_ (Immunohistochemistry)

| **Organ** | **Group** | **6 hpi^s^** | **12 hpi^s^** | **24hpi^s^** | **45-47hpi^d^** | **48hpi^s^** | **53-132hpi^d^** | **14 dpi^s^** | **Cells expressing viral antigen** |
| --- | --- | --- | --- | --- | --- | --- | --- | --- | --- |
| Nasal cavity | 10^2^ EID_50_ | - | - | - | a | - | a | - | Chondrocytes, epithelial cells, mononuclear cells, and endothelial cells |
|  | 10^4^EID_50_ | - | - | - | a | ++ | ++ | - |  |
|  | 10^6^EID_50_ | - | - | ++ | ++ | b | b | b |  |
| Trachea | 10^2^ EID_50_ | - | - | - | a | - | a | - | Chondrocytes and epithelial cells |
|  | 10^4^EID_50_ | - | - | - | a | ++ | ++ | - |  |
|  | 10^6^EID_50_ | - | - | + | ++ | b | b | b |  |
| Lungs | 10^2^ EID_50_ | - | - | - | a | - | a | - | Mononuclear cells and endothelial cells |
|  | 10^4^EID_50_ | - | - | - | a | +++ | +++ | - |  |
|  | 10^6^EID_50_ | - | - | + | ++ | b | b | b |  |
| Brain | 10^2^ EID_50_ | - | - | - | a | - | a | - | Glial cells, ependymal cells, neurons, and granular cells of cerebellum |
|  | 10^4^EID_50_ | - | - | - | a | ++ | ++ | - |  |
|  | 10^6^EID_50_ | - | - | - | ++ | b | b | b |  |
| Spleen | 10^2^ EID_50_ | - | - | - | a | - | a | - | Mononuclear cells |
|  | 10^4^EID_50_ | - | - | - | a | +++ | +++ | - |  |
|  | 10^6^EID_50_ | - | - | + | +++ | b | b | b |  |
| Heart | 10^2^ EID_50_ | - | - | - | a | - | a | - | Cardiac myocytes |
|  | 10^4^EID_50_ | - | - | - | a | +++ | +++ | - |  |
|  | 10^6^EID_50_ | - | - | + | +++ | b | b | b |  |
| Liver | 10^2^ EID_50_ |  |  | - | a | - | a | - | Kupffer cells and hepatocytes |
|  | 10^4^EID_50_ | - | - | - | a | ++ | ++ | - |  |
|  | 10^6^EID_50_ | - | - | + | ++ | b | b | b |  |
| Kidney | 10^2^ EID_50_ | - | - | - | a | - | a | - | No staining |
|  | 10^4^EID_50_ | - | - | - | a | - | - | - |  |
|  | 10^6^EID_50_ | - | - | - | - | b | b | b |  |
| Intestine | 10^2^ EID_50_ | - | - | - | a | - | a | - | Mononuclear cells |
|  | 10^4^EID_50_ | - | - | - | a | - | + | - |  |
|  | 10^6^EID_50_ | - | - | - | + | b | b | b |  |
| Pancreas | 10^2^ EID_50_ | - | - | - | a | - | a | - | Pancreatic acinar cells |
|  | 10^4^EID_50_ | - | - | - | a | - | + | - |  |
|  | 10^6^EID_50_ | - | - | - | + | b | b | b |  |
| Thymus | 10^2^ EID_50_ | - | - | - | a | - | a | - | Mononuclear cells |
|  | 10^4^EID_50_ | - | - | - | a | - | ++ | - |  |
|  | 10^6^EID_50_ | - | - | - | ++ | b | b | b |  |

* - = None; + = Infrequent staining, ++ = Frequent staining, and +++ = Diffuse staining; a - No birds were euthanized or died; b - All birds died before 48 hpi in 10^6^ EID_50_ group; s - Birds were euthanized at the respective interval; d - Dead birds
